# A human embryonic limb cell atlas resolved in space and time

**DOI:** 10.1101/2022.04.27.489800

**Authors:** Bao Zhang, Peng He, John E Lawrence, Shuaiyu Wang, Elizabeth Tuck, Brian A Williams, Kenny Roberts, Vitalii Kleshchevnikov, Lira Mamanova, Liam Bolt, Krzysztof Polanski, Rasa Elmentaite, Eirini S Fasouli, Martin Prete, Xiaoling He, Nadav Yayon, Yixi Fu, Hao Yang, Chen Liang, Hui Zhang, Raphael Blain, Alain Chedotal, David R. FitzPatrick, Helen Firth, Andrew Dean, John C Marioni, Roger A Barker, Mekayla A Storer, Barbara J Wold, Hongbo Zhang, Sarah A Teichmann

## Abstract

Human limbs emerge during the fourth post-conception week as mesenchymal buds which develop into fully-formed limbs over the subsequent months. Limb development is orchestrated by numerous temporally and spatially restricted gene expression programmes, making congenital alterations in phenotype common. Decades of work with model organisms has outlined the fundamental processes underlying vertebrate limb development, but an in-depth characterisation of this process in humans has yet to be performed. Here we detail the development of the human embryonic limb across space and time, using both single-cell and spatial transcriptomics. We demonstrate extensive diversification of cells, progressing from a restricted number of multipotent progenitors to myriad mature cell states, and identify several novel cell populations, including neural fibroblasts and multiple distinct mesenchymal states. We uncover two waves of human muscle development, each characterised by different cell states regulated by separate gene expression programmes. We identify musculin (MSC) as a key transcriptional repressor maintaining muscle stem cell identity and validate this by performing MSC knock down in human embryonic myoblasts, which results in significant upregulation of late myogenic genes. Through integration of multiple anatomically continuous spatial transcriptomic samples, we spatially map single-cell clusters across a sagittal section of a whole fetal hindlimb. We reveal a clear anatomical segregation between genes linked to brachydactyly and polysyndactyly, and uncover transcriptionally and spatially distinct populations of mesenchyme in the autopod. Finally, we perform scRNA-seq on murine embryonic limbs to facilitate cross-species developmental comparison at single-cell resolution, finding substantial homology between the two species.

## Introduction

Human limb buds emerge by the end of the 4th post conceptional week (PCW4) and develop to form arms and legs during the first trimester. By studying model organisms such as the mouse and chick, it is known that development of the limb bud begins in the form of two major components. The multipotent parietal lateral plate mesodermal (LPM) cells condense into the skeletal system as well as forming tendon, fibrous and smooth muscle populations, whilst skeletal muscle progenitor (SkMP) cells migrate from the paraxial mesoderm to the limb field, forming striated muscle^1,2^. These multipotent progenitors are encapsulated within a thin layer of ectoderm, a subset of which (termed the apical ectodermal ridge/AER) governs mesenchymal cell proliferation and aids in the establishment of the limb axes through fibroblast growth factor (FGF) signalling^3^. The limbs continue to mature in a proximal-distal manner, such that by PCW8 the anatomies of the stylopod, zeugopod and autopod are firmly established. This maturation is tightly controlled by a complex system of temporally and spatially restricted gene expression programmes^4–7^. As with any complex system, small perturbations in even a single programme can result in profound changes to the structure and function of the limb^8^. Indeed, approximately 1 in 500 humans are born with congenital limb malformations^9,10^.

Although model organisms have provided key insights into cell fates and morphogenesis that are translatable to human development and disease, at present it remains unclear how precisely these models recapitulate human development. Furthermore, the lack of complementary spatial information in such studies precludes the assembly of a comprehensive tissue catalogue that provides a global view of human limb development in space and time. Encouragingly, the Human Developmental Cell Atlas community has recently applied cell-atlasing technologies such as single cell and spatial transcriptomics to several tissues to give novel insights into development and disease^11–16^. The application of these techniques to human embryonic and fetal tissue therefore holds much promise in furthering our understanding of the developing human limb^17,18^.

In this study, we performed single-cell transcriptomic sequencing (scRNA-seq) and spatial transcriptomic sequencing to reconstruct an integrated landscape of the human hindlimb (or the lower limb) during first trimester development. We then performed scRNA-seq on murine embryonic limbs in order to compare the process of limb development across species at this level of resolution. Our results detail the development of the human limb in space and time at high resolution and genomic breadth, identifying 67 distinct cell clusters from 125,955 captured single cells, and spatially mapping these across four timepoints to shed new light on the dynamic process of limb maturation. In addition, our spatial transcriptomics data gives insights into the key patterning and morphogenic pathways in the developing limb, with a focus on genes associated with limb malformation.

Finally, our integrated analysis of human and murine limb development across corresponding time periods reveals extensive homology between a classical model organism and the human, underlining the importance and utility of such models in understanding human disease and development. Our study provides a unique resource for the developmental biology community, and can be freely accessed at https://limb-dev.cellgeni.sanger.ac.uk/.

## Results

### Cellular heterogeneity of the developing limb in space and time

To track the contribution of the different lineages in the developing limb, we collected single-cell embryonic limb profiles from PCW5 to PCW9 (Fig. 1a). This time window covers early limb forming stages as well as later stages of limb maturation. In total, we analysed 125,955 single-cells that passed quality control filters (Extended Data Fig. 1a). After cell cycle expression module removal by regression, and batch correction (see Methods), we identified 67 cell clusters (Fig. 1b; see Methods; Extended Data Fig. 1b and Extended Data Table 1 for marker genes).

**Figure 1.**
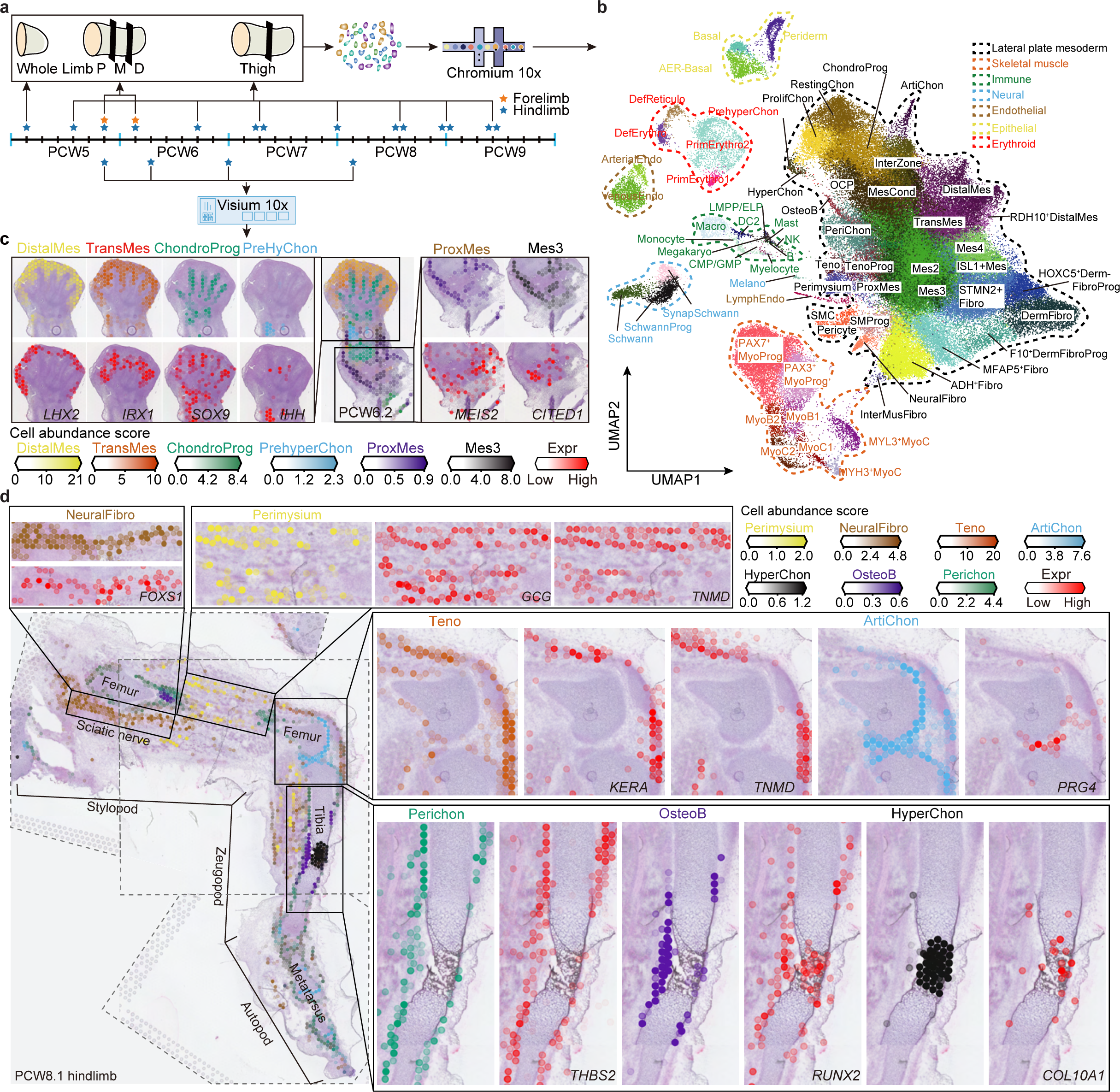
A single-cell temporal-spatial atlas of human embryonic limb. **a**, Overview of human embryonic developmental time points sampled and experimental scheme. The asterisk marks the sampling time point. P, proximal; M, middle; D, distal; PCW, post conceptional week. **b**, Uniform manifold approximation and projection (UMAP) visualization of 125,955 human embryonic limb cells. Sixty-seven cell clusters from seven lineages are labeled in the UMAP. Mes, mesenchyme; ProxMes, proximal Mes; TransMes, transitional Mes; MesCond, mesenchymal condensate cell; OCP, osteochondral progenitor; ChondroProg, chondrogenic progenitor; Chon, chondrocyte; ProlifChon, proliferating Chon; PrehyChon, prehypertrophic Chon; HyperChon, hypertrophic Chon; PeriChon, perichondrium; OsteoB, osteoblast; ArtiChon, articular Chon; Teno, tenocyte; TenoProg, Teno progenitor; MyoProg, myogenic progenitor; MyoB, myoblast; MyoC, myocyte; Fibro, fibroblast; InterMusFibro, muscle interstitial Fibro; DermFibro, dermal Fibro; DermFibroProg, DermFibro progenitor; SMProg, smooth muscle progenitor; SMC, smooth muscle cell; SchwannProg, Schwann progenitor; SynapSchwann, synaptic Schwann; Melano, melanocyte; AER, apical ectodermal ridge; Endo, endothelial; ArterialEndo, arterial Endo; VenousEndo, venous Endo; LympEndo, lymphatic Endo; LMPP/ELP, lymphoid-primed multipotent progenitor/early lymphoid progenitors; CMP/GMP, common myeloid progenitors/granulocyte-monocyte progenitors; NK, Natural killer; DC2, Dendritic Cell 2; Macro, macrophage; Megakaryo, megakaryocyte; DefErythro, definitive erythrocyte; DefReticulo definitive reticulocyte; PrimErythro, primitive erythrocyte. **c-d**, Spatially resolved heatmaps across tissue sections from the PCW6.2 (c) and PCW8.1 (d) human hind limbs showing spatial distribution of selected cell clusters and corresponding marker genes.

34 of these cell clusters represent cells derived from the LPM. They contain mesenchymal, chondrocyte, osteoblast, fibroblast and smooth muscle cell states involved in the maturation of cartilage, bone and other connective tissues, consistent with previous investigations of the cellular makeup of the limb^19^. In addition to these cells, a further eight states form a complete lineage of muscle cells that migrate as PAX3+ progenitors from the somite. These go on to differentiate in the limb to form myoprogenitors and myotubes.

Other non-LPM cell clusters include four of the primitive and definitive erythrocytes, ten clusters of immune cells, three clusters of vascular endothelial cells, three clusters of glial cells and a single melanocyte cluster. Finally, we identified three epithelial cell clusters (Extended Data Fig. 2a), among which was a cluster broadly expressing SP8, and WNT6 (Extended Data Fig. 2b), likely representing distal limb ectodermal cells. Indeed, a small number of these cells (n=9), originating from PCW5 & 6, strongly expressed FGF8; a gene characteristic of the AER (Extended Data Fig. 2c, d). We found the expression of FGF8 to be similarly constrained at PCW5 using RNA in-situ hybridisation (RNA-ISH) (Extended Data Fig. 2e).

Examining the relative abundance of each of the major cell states across different developmental ages revealed how the cellular composition of the developing limb changes over time. Within each of the aforementioned lineages, a clear pattern emerged whereby progenitor states were chiefly isolated from PCW5 & 6, with more differentiated cell states emerging thereafter (Extended Data Fig. 3a,b).

To further dissect the cellular heterogeneity with spatial context, and to build on limb patterning principles established in model organisms, we performed spatial transcriptomic experiments for limb samples from PCW5 to PCW8. Using the 10X Genomics Visium chips, we were able to generate transcriptome profiles capturing on average between 1,000 and 5,000 genes per voxel (Extended Data Fig. 1c). We then applied the cell2location package^20^ leveraging cell cluster signatures from our single-cell atlas to deconvolute Visium voxels (See Methods; Extended Data Fig. 1d for QC metrics). The resulting cell composition map of Visium slides at each time point demarcated the tissue section into distinct histological regions (Fig. 1c, d). In the PCW5.6 samples, differing spatial distributions were observed for distal progenitor cell clusters, dividing them into three populations that we name “distal” (LHX2^+^ MSX1^+^ SP9^+^), “RDH10^+^distal” (RDH10^+^ LHX2^+^ MSX1^+^) and “transitional mesenchyme” (IRX1^+^ MSX1^+^) (Extended Data Fig. 4a-f). The distal mesenchymal cells are located at the distal periphery of the limb. Proximal to it are the transitional mesenchyme together with SOX9-expressing chondrocyte progenitors of the developing autopod (Fig. 1c). This novel spatial distinction was accompanied by subtle transcriptomic differences, with the distal mesenchyme expressing a number of genes implicated in digit patterning, including LHX2, IRX1 and TFAP2B (Fig. 1c; Extended Data Fig. 4a-e). Mutations in the latter cause Char syndrome, a feature of which is postaxial polydactyly^21^. The RDH10^+^ distal mesenchyme strongly expresses RDH10, the primary enzyme of retinaldehyde synthesis which is critical in interdigital cell death^22^ (Extended Data Fig. 4e, f). The transitional mesenchyme expresses IRX1/2, key genes in digit formation that establish the boundary between chondrogenic and non-chondrogenic tissue^23,24^ (Fig. 1c; Extended Data Fig. 4d,e). We further examined the precise spatial distributions of these marker genes in the embryonic limb at PCW5 and PCW6 in three dimensions using tissue clearing and light sheet fluorescence microscopy (LSFM), giving further insight into the arrangement of these tissues during development (Supplementary Video 1).

In addition to mesenchyme, prehypertrophic chondrocytes (PHC) expressing Indian Hedgehog (IHH) localised to the mid-diaphysis of the forming tibia and the metatarsals (Fig. 1c). At the proximal limit of the sample, both MEIS2-expressing proximal mesenchymal cells (ProxMes) and CITED1^+^ mesenchymal cells (Mes3) were observed, in keeping with their role in early limb development (Fig. 1c; Extended Data Fig 4e).

For the PCW8 sample, we obtained three anatomically continuous sections from the hindlimb and placed each on separate capture areas of the same Visium chip. We subsequently integrated data from this chip in order to obtain a spatial transcriptomic readout of a complete sagittal section of the hindlimb (Fig. 1d; Extended Data Fig. 5a). At this stage, articular chondrocytes were located at the articular surfaces of the developing knee, ankle, metatarso-phalangeal and interphalangeal joints, while osteoblasts closely matched to the mid-diaphyseal bone collar of the tibia and femur. The perichondrial cells from which they differentiate matched to a comparable region, though they extended along the full length of the tibia and femur (Fig. 1d); a finding confirmed by immunofluorescence staining for RUNX2 and THBS2 alongside the chondrocyte marker COL2A1 (Extended Data Fig. 5b).

Hypertrophic chondrocytes (HCC) expressing collagen X (COL10A1) mapped to the mid-diaphysis of the tibia (Fig.1d). Additionally, we were able to capture glial cells expressing myelin genes (Extended Data Fig. 1b), and an accompanying FOXS1-expressing fibroblast subtype (named “neural fibroblast” by us here) enriched in the periphery of the sciatic nerve in the posterior compartment of the thigh and its tibial division in the deep posterior compartment of the leg (Fig. 1d, Extended Data Fig. 5c-e). We captured only a very small number of neurons (n=28) in our single-cell data, most likely due to the distant location of their cell bodies within the spinal ganglia.

Interestingly, cell states with related (but not identical) transcriptomic profiles did not necessarily occupy the same location, which we are able to quantify based on our cell2location deconvolution analysis of Visium and scRNAseq data. One example is two groups of fibroblasts within the fibroblast lineage. One group of three clusters were co-located with KRT15-expressing basal cells and SFN-expressing cells of the periderm^25^, suggesting a role in the dermal lineage and prompting their annotation as dermal fibroblasts (DermFiB) and their precursors (F10^+^DermFiBP & HOXC5^+^DermFiBP)(Extended Data Fig. 2f-g). While another group of two fibroblast clusters expressing ADH family members (ADH^+^Fibro, InterMusFibro) co-localised with muscle cells, with no presence in the dermal region (Extended Data Fig. 2h-j).

Similarly, we were able to spatially resolve two clusters with subtle transcriptomic differences within the tenocyte lineage. Both clusters expressed the classical tendon markers scleraxis (SCX) and tenomodulin (TNMD), with one population of cells expressing increased biglycan (BGN) and Keratocan (KERA); molecules which play a role in the organisation of the extracellular matrix (ECM), while the other population expressed higher levels of pro-glucagon (GCG) that is important in metabolism (Extended Data Fig. 5f,g). Analysis with cell2location matched the former cluster to the long flexor tendons of the foot, as well as the hamstrings, quadriceps & patellar tendons around the knee joint. The latter cluster, however, matched to the perimysium, the sheath of connective tissue that surrounds a bundle of muscle fibres (Fig. 1d; Extended Data Fig. 5h). We therefore annotated these clusters as tenocyte (Teno) and perimysium, respectively.

Overall, these findings provide new insights into the subtle transcriptomic differences within cell compartments including the muscle, tendon, bone and stromal lineages. This integrated analysis serves as an example of how spatial transcriptomic methodologies can improve our understanding of tissue architecture and locate cell states themselves within the context of developmental dynamics of an anatomical structure such as the whole limb.

### Patterning, morphogenesis and developmental disorders in the limb

During organogenesis of the limb, individual cell identities are in part, determined by their relative position within the limb. This developmental patterning is controlled by a complex system of temporally and spatially restricted gene expression programmes. For example, key aspects of proximal-distal patterning are controlled by the AER^26,27^. In contrast anterior-posterior axis specification is chiefly controlled by the zone of polarising activity (ZPA) through SHH signalling^28–30^. Within the autopod, precise regulation of digit formation is mediated through interdigital tissue apoptosis^31–33^. We utilised Visium spatial transcriptomic data to explore the locations of transcripts of all these classic pattern-forming genes on the same tissue section, finding consistency with classical expression patterns first identified in model organisms (Extended Data Fig. 6a-e). This included several key genes known to govern proximal identity, including MEIS1 & 2, PBX1 and IRX3^34–37^, as well as genes regulating limb outgrowth and distal morphogenesis such as WNT5A, GREM1, ETV4 and SALL1^38–41^. Similarly, classical mammalian anterior-posterior (AP) genes were captured, including HAND1, PAX9, ALX4 and ZIC3 (anterior) and HAND2, SHH, PTCH1 and GLI1 (posterior)^28,42–48^.

The homeobox (HOX) genes are a group of 39 genes split into four groups termed “clusters”, each of which is located on a separate chromosome. During limb development, genes in the A & D clusters act in concert with the aforementioned axis-determining genes to dictate limb patterning in mammals^49^. In mice, these genes are expressed in two waves. The first wave occurs in the nascent limb bud (prior to the period captured in our samples) with expression progressively restricted to its posterior margin with increasing 5’ position. During the second wave, this expression pattern is no longer seen in the A cluster, but persists in the 5’ members of the D cluster^50,51^. Our spatial transcriptomic data captured the expression patterns of the A and D clusters at PCW5.6 (Extended Data Fig. 6f). As expected, their expression matches the second wave of expression in mice, with a loss of asymmetry in the HOXA cluster and its maintenance in the HOXD cluster. For both clusters, an increase in group number corresponded to more distally restricted expression, with group 13 genes limited to the most distal part of the autopod. An exception to this was HOXA11 expression, which showed no overlap with HOXA13, in keeping with the expression pattern of these two genes during the second HOX wave in mice. In fact, our data revealed a clear switch to the antisense transcript of HOXA11 in the distal limb (Extended Data Fig. 6f). This mutual exclusivity in expression domain is thought to be due to HOXA13/D13-dependent activation of an enhancer that drives antisense transcription of HOXA11 in pentadactyl limbs^52^

In order to investigate gene expression patterns during digit formation, we obtained coronal sections through a PCW6.2 foot plate to reveal the five forming digits together with the intervening interdigital space (IDS; Fig. 2a). We then performed louvain clustering on these slides to unbiasedly annotate digital, interdigital, distal mesenchyme and other regions (Fig. 2a, see Methods). Differential expression testing between digital and interdigital regions across two adjacent sections demonstrated an enrichment of classical genes involved in interdigital cell death in the IDS, such as BMP7, BMP2 and ADAMTS1^53,54^ (Fig. 2b). Interdigital regions also showed an enrichment of retinol dehydrogenase 10 (RDH10) and Cellular Retinoic Acid Binding Protein (CRABP1), whereas the retinoic acid (RA) metabolising enzyme CYP26B1 was upregulated in the digital regions (Fig. 2b). We further validated their spatial patterns at single-cell level using RNA-ISH (Fig. 2c). These findings underline the importance of RA in triggering interdigital cell death in the hand and foot plate^32,55^. Other genes identified as digit-specific included TGFB2, a vital molecule in interphalangeal joint specification, and the regulator of chondrogenesis, WWP2 ^56,57^. In addition, PIEZO2, which promotes bone formation via calcium-dependent activation of NFATc1, YAP1 and ß-catenin, was restricted to the digits, together with the calcium-binding molecule C1QL1, which has been shown to correlate with COL2A1 expression during in vitro chondrogenesis ^58,59^(Fig. 2b).

**Figure 2.**
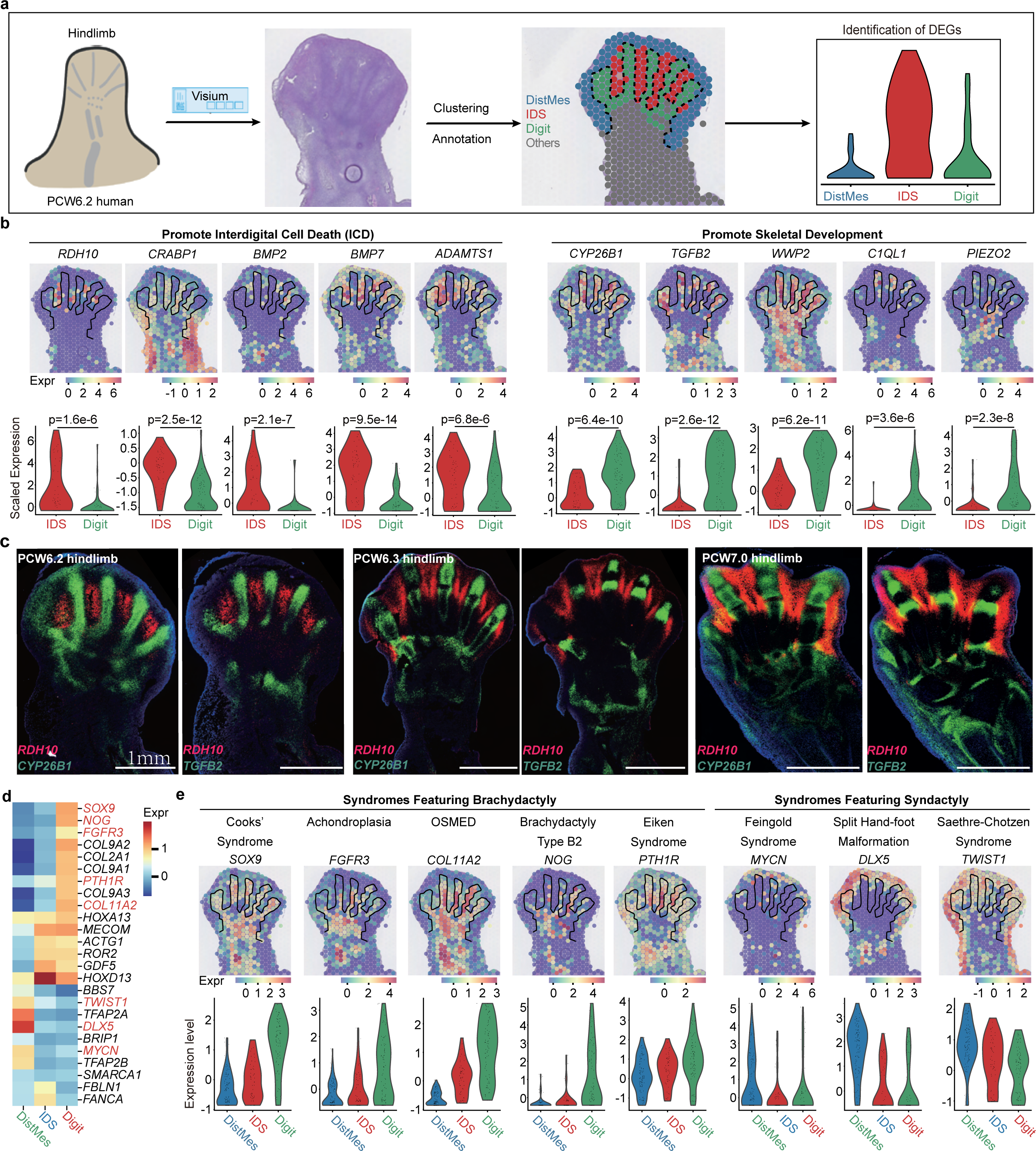
Spatial expression pattern of genes involved in digit formation and phenotype. **a**, Overview of experimental scheme to identify genes involved in digit formation and interdigital cell death (ICD). IDS, interdigital space. DistMes, distal mesenchyme **b**, Spatially resolved heatmaps across tissue sections from the PCW 6.2 human lower limb showing spatial expression pattern of genes promoting ICD (left panel) and digital tissue survival (right panel), and violin plots showing significance of differences between genes of digital region and IDS. The color bar indicates the normalized and log-transformed expression level. **c**, RNA-ISH of tissue sections from human hind limb showing the expression pattern of RDH10, CYP26B1 and TGFB2. Scale bar, 1mm. **d**, Heatmap showing the expression differential patterns of genes associated with digit malformation. The color bar indicates the Z-score of expression level. **e**, Spatially resolved heatmaps across tissue sections from the PCW 6.2 human hind limb showing spatial expression pattern of selected genes associated with digit malformation, as well as violin plot showing the differences in genes between digital region, IDS and distal mesenchyme (DistMes). The color bar indicates the normalized and log-transformed expression level.

Finally, we histologically annotated each digit in the PCW6.2 foot plate to search for genes that vary with digit identity (Extended Data Fig. 6i). We identified four genes that were upregulated in the great toe including ID2 and ZNF503, both of which are known to have anterior expression domains in the limb, as well as the regulator of cell proliferation PLK2 and the cancer-associated gene LEMD-1^60–63^. HOXD11 was downregulated in the great toe, in keeping with its posterior prevalence. We found no differentially expressed genes in the remaining digits, though statistical power was limited by the small number of voxels occupied by each.

We next cross-referenced the list of spatially differentially expressed genes against a list of 2300 single gene health conditions. We found genes involved in several types of isolated (or non-syndromic) brachydactyly (BD) were significantly upregulated in the digital tissue (Fig. 2d,e; full list of DE genes shown in Extended Data Table 2). These included NOG (brachydactyly type B2), PTH1R (Eiken syndrome), COL11A2 (Oto-Spondylo-Mega-Epiphyseal Dysplasia / OSMED), SOX9 (Cook’s syndrome) and FGFR3 (Achondroplasia)^64,65,66^. Conversely, genes which are varied in syndromes with syndactyly as part of their phenotype were significantly upregulated in the IDS and distal mesenchyme. These include DLX5 (split hand-foot malformation), MYCN (Feingold Syndrome type 1), and TWIST1 (Saethre-Chotzen Syndrome)^67–70^. Where murine models of the aforementioned heritable conditions exist, their phenotype is broadly comparable to the human (Extended Data Table 3)^71,72^ ^65,66,69,73–87^. Thus, our spatial atlas provides a valuable reference of gene expression under homeostatic conditions for comparison with genetic variations for which phenotypes may begin to penetrate during embryonic development.

### Regulation of cell fate decisions of mesenchymal-derived lineages

Our single-cell and spatial atlases revealed a high diversity of mesenchymal-derived cell clusters. In order to better understand what transcriptional mechanisms may control their specification, we inferred cell-fate trajectories in the 34 mesenchyme-associated states by combining diffusion maps, partition-based graph abstraction (PAGA) and force-directed graph (FDG) (see methods).

As expected, the global embedding resembled a ‘spoke-hub’ system, whereby multipotent mesenchymal cells are embedded centrally, with clusters of lineage-committed cells radiating outward as they begin to express classical cell-type specific marker genes (Fig. 3a, b). The central hub of mesenchymal cells consisted of six clusters with subtle differences in their transcriptome (Extended Data Fig. 4e). A first population, proximal mesenchymal cells (ProxMes) expressed the regulator of proximal (stylopod) identity, MEIS2, together with WT1, which marks the point of limb-torso junction and plays an unknown role in limb development ^88^(Fig. 3a,b; Extended Data Fig. 4e, g). A similar population, here named Mesenchyme 1 (Mes1), exhibited a similar expression profile but lacked WT1 (Fig. 3a,b). This likely represents mesenchyme in the proximal limb just distal to the limb-torso junction (Extended Data Fig. 4e, g). We also identified a population of mesenchymal cells that expressed ISL1 (ISL1^+^Mes) in addition to MEIS2, but not WT1 (Extended Data Fig. 4e,g). These cells represent a mesenchymal niche within the posterior aspect of the developing hindlimb^89^. Two further clusters (Mes2, Mes3) of mesenchyme expressed CITED1, a molecule which localises to the proximal domain of the limb and plays an unclear role in limb development^90^(Extended Data Fig. 4e, g). Mes3 strongly expressed CITED1, whereas Mes2 exhibited lower expression, and co-expressed MEIS2, suggesting proximal-anterior location in the limb^91^ (Extended Data Fig. 4e, g).The Mes4 cluster exhibited similar overall expression patterns to distal and transitional mesenchymal cells though lacked LHX2 and IRX1, expressing low levels of PRAC1, a molecule identified as maintaining a prostate gland stem cell niche but with no known role in limb development^92^.

**Figure 3.**
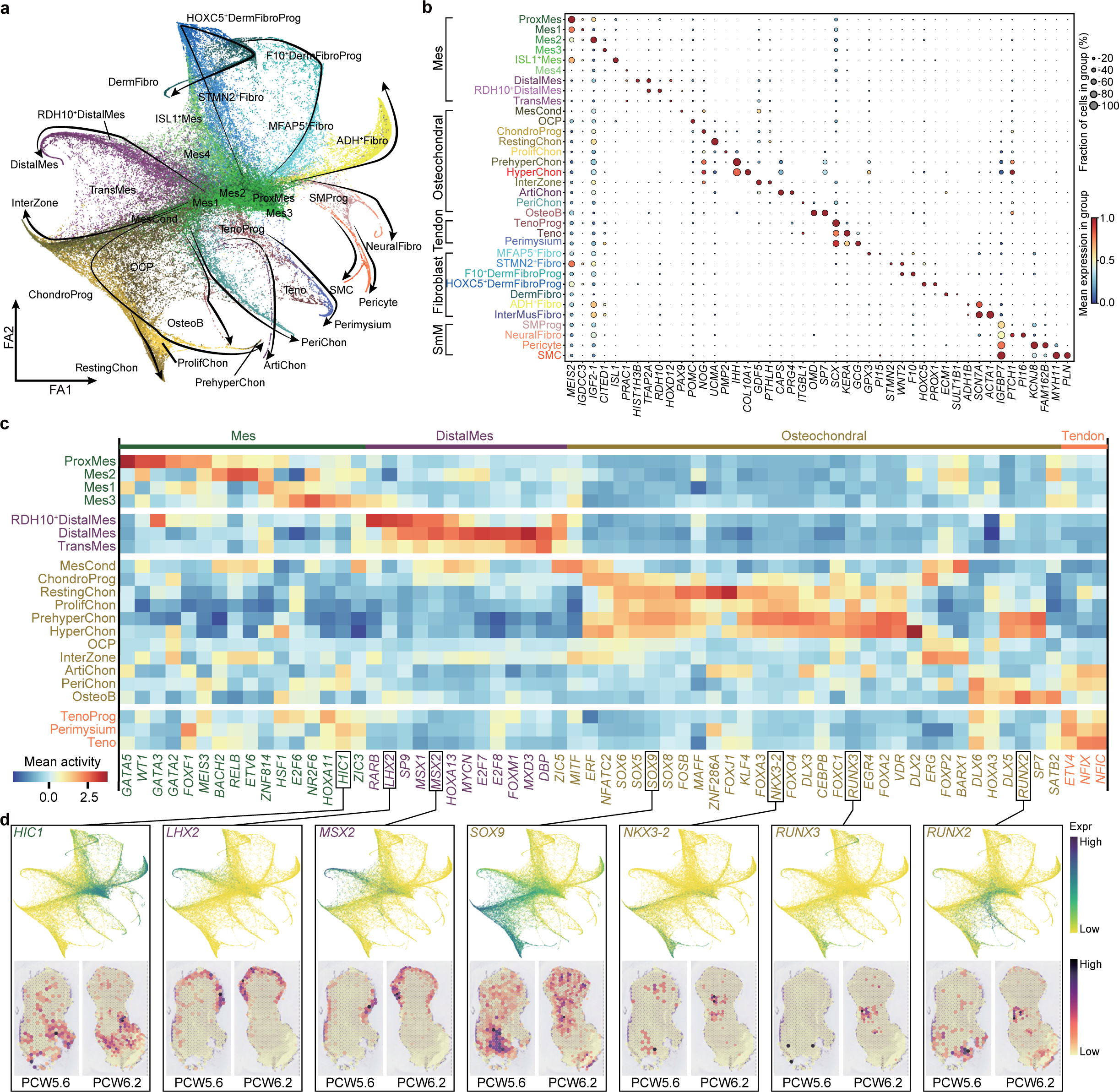
Cell lineage diversification and transcription factor (TFs) specificity of lateral plate mesoderm (LPM) during human embryonic limb development. **a**, Force-directed graph layout of cells associated with the LPM, colored by cell clusters. Black arrows indicate the direction of cell differentiation. Cluster abbreviations same as Fig. 1. **b**, Dot plot showing selected marker genes for each cell cluster. The color bar indicates the average expression level in linearly scaled values. **c**, Heatmap illustrating the vertically normalized mean activity of selected TFs for each cell cluster. **d**, Force-directed graph (top) and spatially resolved heatmaps across tissue sections from human lower limb (bottom) showing expression pattern of selected TFs. PCW, post conception week.

Examining the abundance of different clusters by developmental age revealed how cellular heterogeneity within the mesenchymal compartment evolves during limb development (Extended Data Fig. 7c). During PCW5, the majority of the cells captured were mesenchymal progenitors. This was particularly notable at PCW5.1 and 5.4, where mesenchyme accounted for 85% and 65% of all cells respectively. The relative abundance of mesenchymal progenitor cells in the limb declined thereafter, with almost none present at PCW8 & 9.

We next performed transcription factor (TF) network inference using the SCENIC package to identify distinct modules of active TF networks associated with progression through each lineage^93^. A multitude of TF networks were predicted to be active in these progenitor populations (Fig. 3c, d; Extended Data Fig. 7a, b; Extended Data Table 4). In addition to WT1 and MEIS family members, several GATA factors were predicted to be active in the proximal mesenchyme. This included GATA5, which was recently identified as a putative proximo-distal patterning gene in the Xenopus limb, and GATA3, which has been detected in the proximal developing mouse limb ^94,95^. HOXA11, which defines the zeugopod, was active in Mes3, in keeping with the aforementioned lack of MEIS1/2 expression in this cluster. Cells of the distal mesenchyme showed activation of a distinct module of TFs, including LHX2 and MSX1/2, as previously described in the mouse, as well as HOXA13, which defines the autopod^96–98^. Interestingly, HIC1 was predicted to have low activity in several mesenchymal populations, with high activity in Mes3 (Fig. 3c, d). HIC1^+^ mesenchymal cells are known to migrate into the limb from the hypaxial somite, eventually differentiating into a range of tissues such as chondrocytes and tenocytes, whilst maintaining HIC1 expression ^99^. Indeed, some chondrocyte populations and all tendon populations showed HIC1 activity (Fig. 3c, d).

The chondrocyte lineage increased in number steadily over time, accounting for 25% of the cells captured at PCW5.6, increasing to 50% at PCW7.2. Within this lineage, a shift from progenitors to more mature clusters was observed during the period studied (Extended Data Fig. 3a, b). Mesenchymal condensate cells, SOX^low^/COL2A1^low^/PRRX1^hi^ osteochondral progenitors (OCP) and immature, SOX9^hi^ / COL2A1^low^ chondrocyte progenitors giving way to three populations of SOX9^hi^/COL2A1^hi^ chondrocytes: resting (expressing UCMA), proliferating (with a greater proportion of cells in G2/M/S phases, but lacking UCMA or IHH) and prehypertrophic chondrocytes (expressing IHH) (Fig. 3b,c; Extended Data Fig. 7d-g; Extended Data Table1). In addition, our single cell data captured a small number (n=14) of hypertrophic chondrocytes expressing COL10A1 and MMP13 (Extended data Fig. 7d). Curiously, both PAGA and RNA velocity analysis suggested chondrocyte progenitors for an individual sample may progress to either the resting state prior to proliferation or proceed directly to proliferation (Fig. 3a; Extended data Fig. 7e, f). However these computational predictions should be interpreted cautiously, and further work is required to investigate this finding.

Progression through the chondrocyte lineage was accompanied by changes in regulon activity (Fig. 3c, d). For example, the transition from mesenchymal condensate to committed chondrocytes was associated with activity in the master regulators of chondrogenesis - SOX5, 6 and 9, with the latter localising to chondrocyte condensations at PCW 5.6 and the developing tibia, fibula and digits at PCW6.2 (Fig. 3d)^100^. This trend was observed for other known regulators of chondrogenesis, including MAFF and NKX3-2^101,102^. Interestingly, FOXJ1 was predicted to have similar activity in the chondrocyte lineage, particularly in resting chondrocytes. In addition to its established role in ciliation, this TF has been shown to regulate dental enamel development ^103^. Furthermore, like SOX5 and 9, chondrogenesis is regulated by IRX1; a TF which specifies the digits and establishes the boundary between chondrogenic and non-chondrogenic tissue in the developing chick limb^23^. Several known regulators of chondrocyte hypertrophy were active in PHCs and HCCs, including Osterix (SP7), DLX2/3 and RUNX3, with the latter localising to the tibial diaphysis at PCW 6.2^104–106^. RUNX2 was, as expected, predicted to be active in osteoblasts and the perichondrial cells from which they are derived, in addition to PHCs and HCs, with expression localising to the tibial diaphysis (Fig. 1d). Finally, the osteogenic regulator SATB2 was highly specific to osteoblasts ^107^.

In order to capture the cells of the interzone, mesenchymal cells that reside at the sites of future synovial joints and give rise to their constituent parts, we sectioned four limbs (two forelimbs, two hindlimbs) into proximal, middle (containing the knee / elbow interzone regions) and distal segments. Our data indeed contained a cluster expressing the classical interzone marker GDF5 and which gave rise to articular chondrocytes expressing lubricin (PRG4; logFC=5.15, p=4.1E-26) (Fig. 3b; Extended Data Fig. 7d). Intriguingly, the articular chondrocytes were not predicted to exhibit SOX5/6/9 activity, but instead showed activity in FOXP2, a negative regulator of endochondral ossification, and ERG, which directs chondrocytes down a permanent developmental path within the joint^108,109^ (Fig. 3c). The role of SOX9 in articular cartilage development and homeostasis is uncertain, with inducible loss in mice resulting in no degenerative change at the joint postnatally^110,111^.

Tendon progenitors expressing high levels of SCX but low TNMD emerged during PCW5 before declining in number and being replaced by tenocytes and perimysium expressing high levels of TNMD from PCW7 onward (Fig. 3b, Extended Data Fig. 7a). Of note, our data captured a significant population of SCX^+^/SOX9^+^ cells; a population previously shown to give rise to cells of the entheses (Extended Data Fig. 7h, i)^112^. Visium spatial transcriptomic assays and RNA in-situ hybridisation confirmed co-expression (Extended Data Fig. 7j, k). Several TFs implicated in tenogenesis were predicted to be active in these clusters, including ETV4 and NFIX^113,114^(Fig. 3c).

Finally, different fibroblast and smooth muscle populations within the limb exhibited clearly distinct TF activities (Extended Data Fig. 7a, b). For example, dermal fibroblasts showed activity in known regulators of this lineage, including RUNX1 and TFAP2C ^115,116^, whereas smooth muscle cells (SMC) and their precursors (SMProg) both showed activity in GATA6, which is thought to regulate their synthetic function ^117^. In addition, SMC showed activity in TFs with known roles in smooth muscle function, such as ARNTL^118^.

### Regulation of embryonic and fetal myogenesis

Limb muscle originates from the dermomyotome in the somite^1,2^. Classically, its formation begins with delamination and migration from the somite regulated by PAX3 and co-regulators such as LBX1 and MEOX2, followed by two subsequent waves of myogenesis: embryonic and fetal^119^. During embryonic myogenesis, a portion of PAX3^+^ embryonic skeletal muscle progenitors are destined to differentiate and fuse into multinucleated myotubes. These primary fibres act as the scaffold for the formation of secondary fibres derived from PAX7^+^ fetal skeletal muscle progenitors, which are themselves derived from PAX3^+^ muscle progenitors^120–122^.

To dissect these limb muscle developmental trajectories in detail from our human data, we took cells from the eight muscle states, re-embedded them using diffusion mapping combined with PAGA and FDG. Three distinct trajectories with an origin in PAX3^+^ skeletal muscle progenitors (PAX3^+^ SkMP) emerged (Fig. 4a). The first trajectory (labelled 1st Myogenesis) starts from PAX3^+^ SkMP and progresses through an embryonic myoblast state (MyoB1) followed by an early embryonic myocyte state (MyoC1), and finally arrives at mature embryonic myocytes. This trajectory is in keeping with embryonic myogenesis. Along the second trajectory, the PAX3^+^ SkMP leads to PAX3^+^PAX7^+^ cells, followed by a heterogeneous pool of PAX7^+^ SkMP cells that are mostly MyoD negative (Fig. 4a,b). This represents a developmental path that generates progenitors for subsequent muscle formation and regeneration. The final trajectory (labelled 2nd Myogenesis) connects cell states that express PAX7 first to fetal myoblasts (MyoB2), then early fetal myocytes (MyoC2) and finally mature fetal myocytes (Fig. 4a,c).

**Figure 4.**
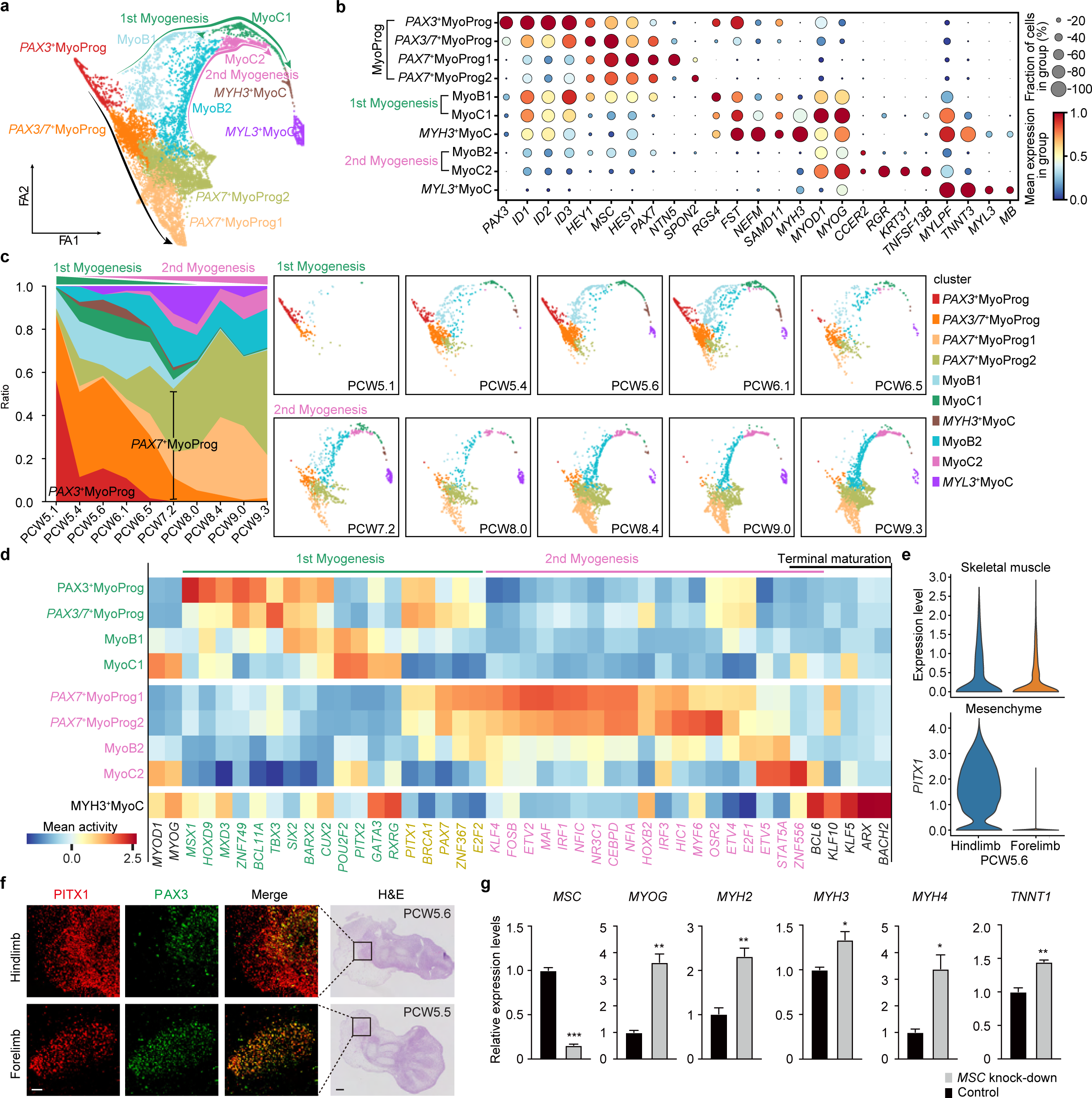
Cell trajectory and transcription factors (TFs) conversion of embryonic and fetal myogenesis during human embryonic limb development. **a**, Force-directed graph layout of cells associated with the myogenesis, coloured by cell clusters. Green and pink arrows indicate the direction of first and second myogenesis, separately. MyoProg, myogenic progenitor; MyoB, myoblast; MyoC, myocyte. **b**, Dot plot showing expression pattern of selected marker genes for each cell cluster. The colour bar indicates the average expression level in linearly scaled values. **c**, Fraction of cell type per time point (left) and Force-directed graphs layout of cells from each time point, coloured by cell clusters. **d**, Heatmap illustrating the vertically normalised mean activity of filtered TFs for each cell cluster. **e**, Violin plot showing the expression level of PITX1 between human fore-and hind-limb. **f**, Immunofluorescence co-staining (scale bar: 50 *μ*m) of PITX1 and PAX3 on hind-(top panel) and fore-(bottom panel) limb sections (scale bar: 200 *μ*m) at PCW5. **g**, RT-qPCR analysis of the fold enrichment of indicated myocyte differentiation genes upon knock-down of MSC in human primary embryonic myoblasts. Data are presented as mean ± SEM. *P < 0.05, **P < 0.01, ***P <0.001 (two-sided Student’s t test).

Comparing these myogenic pathways, we noticed that PAX3 expression is almost absent in the fetal myogenic pathway while it persists to late states along the trajectory of the embryonic myogenic pathway (Fig. 4b). This is consistent with a previous study that captured Pax3^+^ Myog^+^ cells in the mouse limb ^123^. Interestingly, ID2 and ID3 that are known to attenuate myogenic regulatory factors ^124,125^ are also more highly expressed in embryonic myogenesis than fetal, which may imply different upstream regulatory networks. Additional genes such as FST, RGS4, NEFM and SAMD11 were also identified to be marking the first myogenic pathway while TNFSF13B, KRT31 and RGR mark the second (Fig. 4b). In fact, Keratin genes have been found to facilitate sarcomere organisation^126^.

Next, we performed SCENIC analysis to search for transcription factors driving each myogenic stage. A large number of stage-specific transcription factors were identified (Fig. 4d; Extended Data Fig. 8a; Extended Data Table 4). Whilst the master regulators MYOD1 and MYOG showed similar activities across fetal and embryonic myogenesis, several TFs were predicted to have higher activity in one or the other. For example, PITX2 exhibited a higher activity score and abundance during embryonic myogenesis than fetal myogenesis, possibly related to its different regulatory roles^127^. By contrast, Its related family member, PITX1, exhibited comparable activity in both trajectories. Interestingly, while known as a hindlimb-specific transcription factor, PITX1 is expressed in both forelimb and hindlimb muscle cells (Fig. 4e; Extended Data Fig. 8b), including a fraction of PAX3^+^ cells as early as PCW5 (Fig. 4f; Extended Data Fig. 8b), suggesting a potential regulatory role in embryonic myogenesis. Other TFs specific to embryonic myogenesis included MSX1, which maintains the early progenitor pool, the MyoD activator SIX2, and the satellite cell homeodomain factor BARX2 ^128–130^. The fetal myogenic trajectory was associated with several TFs involved in regulating myogenesis, such as E2F2 and MYF6.^131,132^.

Complementary to SCENIC analyses focusing on activators, we also investigated several transcriptional repressors such as MSC (also known as Musculin, ABF-1 or MyoR), TCF21(Capsulin), and families of ID, HES and HEY genes. We observed specific expressions of IDs, HEY1, MSC, and HES1 in PAX7^+^ skeletal progenitors (Fig. 4b). The most prominent repressor, MSC is a bHLH transcription factor that has been shown to inhibit MyoD’s ability to activate myogenesis in 10T1/2 fibroblasts^133^ and rhabdomyosarcoma cells^134^. In addition, in C2C12 murine myoblasts, MSC facilitates Notch’s inhibition of myogenesis (although it appears to exhibit functional redundancy in this role)^135^. To test whether human MSC also plays a role in repressing PAX7^+^ skeletal muscle progenitor maturation^136,137^, we knocked down MSC in primary human embryonic limb myoblasts. RT-qPCR results showed profound upregulation of late myocyte genes (Fig. 4g). This suggests that MSC is key to maintaining limb muscle progenitor identity.

### Spatially resolved microenvironments exhibit distinct patterns of cell-cell communication

In order to investigate communication between clusters of cells, we utilised the CellphoneDB python package to identify stage-specific ligand-receptor interactions by cell type in the developing limb^138,139^. This output was then filtered to reveal signalling pathways between co-located populations of cells, determined in an unbiased way using cell2location factor analysis. Our analysis highlighted the role of the WNT signalling pathway in early limb morphogenesis. As has been found in model organisms, WNT5A exhibited a proximo-distal expression gradient, with expression peaking in the distal mesenchymal populations (Fig. 5a-e). Its receptor FZD10 was expressed in the distal ectoderm of the limb at PCW6 with weak mesenchymal expression, though at this comparatively late stage appears to be no longer restricted to posterior regions, as is reported in early limb development in the mouse and chick ^140,141^(Fig 5a-c). Furthermore, our single cell data revealed high expression of the canonical receptor FZD4 in the mesenchymal condensate; a finding supported by RNA-ISH (Fig.5a, d and e). This gives weight to the suggestion from in vitro studies that FZD4 plays a role in initiating chondrogenesis when a collection of mesenchymal cells reaches a critical mass ^142^.

**Figure 5.**
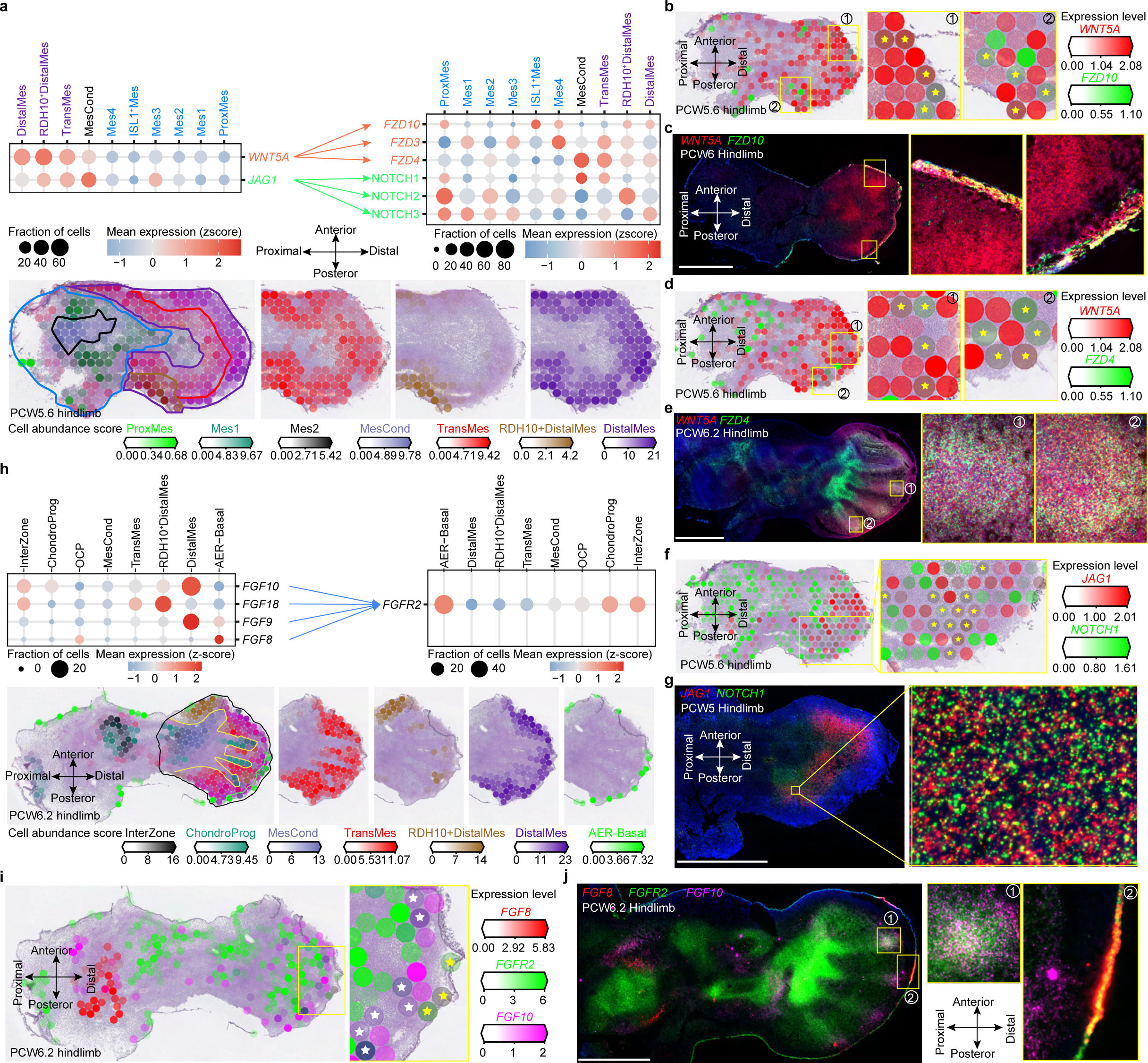
Spatially resolved cell-cell communication. **a**, Dot plots (top panel) showing expression of ligands and cognate receptors in cell clusters. The colour bar indicates the Z-score of average expression level. Mes, mesenchyme; ProxMes, Proximal Mes; MesCond, mesenchymal condensate cell; TransMes, transitional Mes. Spatially resolved heatmaps showing predicted cell-type abundance. **b, d, f**, Spatially resolved heatmaps across tissue sections from PCW5.6 (post conception week 5 plus 6 days) human hindlimb showing spatial expression of WNT5A (b, d), JAG1 (f) and their cognate receptors FZD10 (b), FZD4 (d) and NOTCH1 (f). The colour bar indicates the expression level of normalised and log-transformed counts. **c, e, g**, RNA-ISH of tissue sections from human hind limb showing the expression pattern of WNT5A (c, e), JAG1 (g) and their cognate receptors FZD10 (c), FZD4 (e) and NOTCH1 (g) in situ. Scale bar, 1mm. **h**, Expression dot plots of FGFR2 and its ligands and heatmaps across tissue sections from the PCW 5.6 human hindlimb showing spatially resolved selected mesenchymal cell cluster (separated by colour) signatures. i, Heatmaps across tissue sections from the PCW 6.2 human hindlimb showing spatial expression of FGF8/10 and FGFR2. j, RNA-ISH of FGF8/10 and FGFR2.

In the early (PCW5.6) limb, NOTCH signalling was predicted to occur in its distal posterior aspect through the canonical ligand Jagged (JAG)-1 (Fig. 5a, f). This interaction occurs between adjacent cells, with JAG1 bound to the cell rather than being secreted, triggering proteolytic cleavage of the intracellular domain of NOTCH receptors with varying activity depending on the NOTCH receptor involved^143–145^. JAG1 is induced by SHH in the posterior distal limb;^43,146^ a distribution we confirmed with spatial transcriptomics and show co-localisation at single cell resolution using RNA-ISH (Fig. 5 g). Analysis of single cell data at the corresponding time point showed JAG1 and NOTCH1 to be expressed by several mesenchymal populations within the early limb (Fig. 5a). This finding sheds further light on the mechanisms controlling limb morphogenesis and has implications for conditions where this signalling axis is disrupted, such as the posterior digit absence of Adams-Oliver syndrome and the 5th finger clinodactyly of Alagille syndrome^147,148^.

We captured weak but reproducible signals of FGF8 in the AER epithelial cells across various timepoints while FGF10 was detected in the adjacent distal mesenchyme (Fig. 5h-i). It is known that FGF8 and FGF10 are expressed in the limb ectoderm and distal mesenchyme respectively and form a feedback loop through FGFR2 that is essential for limb induction^149^. Ectodermal FGF8 expression was confirmed via RNA-ISH (Fig. 5j). FGF10 expression was notably restricted in the foot plate next to the distal mesenchyme adjacent to the forming phalanges and excluded in the IDS region and RDH10+ distal mesenchyme(Fig. 5 h,j). This is consistent with expression in lineage tracing experiments in the mouse, where conditional knockdown leads to truncated, webbed digits ^150^. FGF10 has been shown to induce chondrogenesis via FGFR2, which we found to be densely expressed in the chondrocyte progenitors (Fig. 5f). RNA-ISH confirmed this high FGFR2 distribution throughout the skeletal elements of the forming limb (Fig. 5j). The importance of this receptor in skeletal development is highlighted by the limb phenotypes observed in the FGFR2-related craniosynostoses, such as radiohumeral synostosis, arachnodactyly and bowed long bones^151^.

### Homology and divergence between human and murine limb development

Limb development has long been studied in model organisms, while assays directly performed on human samples are less common. To explore differences between mice and humans, we collected 13 mouse limb samples for scRNA-seq, and combined our newly generated data with 18 high-quality (Extended Data Fig. 9a) limb datasets from three previously published studies ^152–154^ to build a comprehensive mouse embryonic limb atlas (Fig. 6a, Extended Data Fig. 9b, c). To compare the mouse and human transcriptome, we used our previously developed alignment algorithm MultiMAP^155^ to align single-cells based on matched orthologs, while also considering information from non-orthologous genes (Fig. 6b). The resulting integrated atlas with aligned cell-type clusters show a highly conserved cell composition between Human and Mouse (Fig. 6b-d; Extended Data Fig. 9d). An independent and separate analysis also shows highly similar developmental trajectories of the skeletal muscle (Extended Data Fig. 8) and LPM-derived lineages (Extended Data Fig. 10), including the finding that PITX1 is expressed in both forelimb and hindlimb muscle cells of both species (Extended Data Fig. 8b).

**Figure 6.**
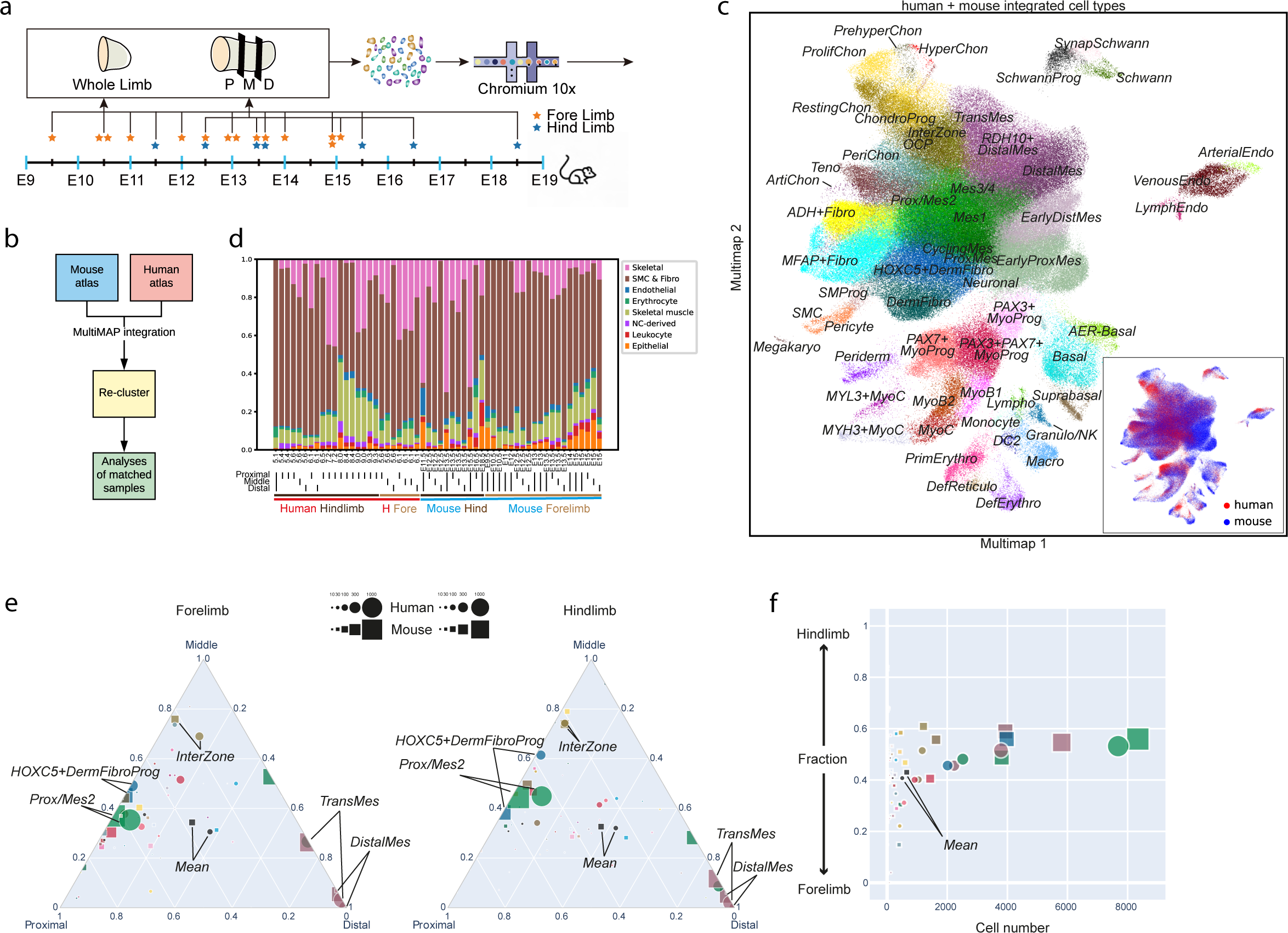
Comparison of single cell atlas between human and mouse limb. **a**, Overview of mouse embryonic developmental time points sampled and experimental scheme. The asterisk marks the sampling time point. P, proximal; M, middle; D, distal. **b**, Overview of analysis pipeline to integrate human and mouse scRNA-seq data. **c**, Multimap layout of integrated cells, coloured by integrated cell-type annotation or species (bottom right). **d**, Broad cell-type proportions of each scRNA-seq library with dissection region, location and species labeled at the bottom. **e-f**, Triangular diagrams (e) showing the cell-type proportion biases towards proximal, middle or distal region of human and mouse forelimb (left panel) and hindlimb (middle panel) and a scatter plot (f) showing the fraction of each cell type’s hindlimb representation. Each cell type or mean is represented by a circle (human) and a square (mouse) with size (square of diameter) meaning average number of cells per segment (proximal/middle/distal).

Some notable differences in the human and mouse datasets were most likely due to sampling differences, such as the greater abundance of PAX3^+^ myoprogenitors in the mouse and the presence of two distinct mesenchymal populations, “Early Distal Mes’’ and “Early Prox Mes” (Extended Data Fig. 8c-e; Extended Data Fig. 10a-b). The vast majority of these cells originate from samples prior to E12; the equivalent developmental stage to the earliest human stage sampled (Extended Data Fig. 8f, Extended Data Fig. 10c). Similarly, the lack of Wt1 expression in the mouse proximal mesenchyme is most likely due to dissection not including the trunk, as this expression pattern was first described in the mouse^88^.

However, we also identified species-specific features when comparing datasets. Mouse limbs contained a higher percentage of epithelial cells and immune cells (Fig. 6d) possibly due to faster maturation of these systems in the mouse. Additionally, in skeletal muscle, a more abundant PAX3+PAX7+MyoProg cell state was observed in humans than in mice. (Extended Data Fig. 8c-e). Whilst the PAX3^+^ pools were highly similar in their gene expression, the mouse data showed curiously low expression of pro myogenic factors Fst and Uchl1, though this may again be due to differences in sample stage from which the PAX3^+^ pools arise^156,157^(Extended Data Fig. 8g). Development in both species involved a heterogenous PAX7^+^ pool, however the degree of heterogeneity appeared greater in the mouse data, with the two clusters (PAX7^+^MyoProg1 & 2) showing clear differential expression of the ECM genes Lamb1, Matn2 and Eln (Extended Data Fig. 8g). The mesenchymal compartments of both species showed substantial similarity in gene expression of different cell clusters (Extended Data Fig 10 d). One interesting finding that held true for both species was the expression of FGF8 in the proximal mesenchyme (Extended Data Fig. 2c-e; Extended Data Fig. 10e, f). This molecule is not classically associated with mesenchymal cell states other than in urodeles, and its role in this human mesenchymal niche is unclear ^158^.

To systematically compare pattern formation between mouse and human limbs, we dissected forelimbs and hindlimbs from a human embryo and a mouse counterpart, each separated into proximal, middle and distal segments to compare with our first trimester human samples (Fig. 1a, Fig. 6a). This allowed us to address the differences between forelimb and hindlimb along the proximo-distal axis at matched time points in human and mouse development.Overall, mice and humans demonstrate highly similar cell cluster compositions along the P-D axis. In both human and mouse forelimbs, proximal mesenchymal cells are enriched towards the proximal end, while distal and transitional mesenchymal cells are highly enriched in the distal part as expected (Fig. 6e). Additionally, interzone cells are enriched in the middle segment, where we intentionally included the joints. The same is true for the hindlimb. Comparison of forelimbs and hindlimbs demonstrated that both humans and mice show minimal differences in terms of cell type composition (Fig. 6f). This suggests that the composition of the developing limb is highly conserved between humans and mice even when pinpointing the broad anatomical regions. To perform a more stringent comparison, we took cells from the thirty-two LPM-derived states to compare ortholog expression signatures between proximal and distal segments in mouse and human. Both species recapitulate known P-D biased genes such as MEIS2 (proximal), LHX2 (distal) and HOX family genes (Extended Data Fig. 9e). Known forelimb/hindlimb biased genes were also captured such as TBX5 specific to the forelimb and TBX4, PITX1 and ISL1 specific to the hindlimb (Extended Data Fig. 9f). Overall, we show that the spatial expression patterns of genes controlling forelimb/hindlimb identity and P-D axis formation are highly similar between mice and humans.

## Discussion

Our developmental limb atlas combines single-cell RNA and spatial transcriptomic analyses of embryonic limb cells from multiple time points in the first trimester in order to form the first detailed characterisation of human limb development across space and time. We identify sixty-seven clusters of cells within eight tissue lineages in the developing limb and place them into anatomical context, building on existing knowledge of cellular heterogeneity gained from model organisms^152^. Our spatial data also reveals the expression of key regulators of limb axis identity in the developing human limb.

In addition to recapitulating model organism biology, our atlas enables the identification of novel cell states. We identify and confirm a population of neural fibroblasts surrounding the sciatic nerve and its tibial division. We also characterise several novel populations of mesenchymal cells as mesenchymal progenitors and distal mesenchyme subtypes that may play unclear roles in limb formation and should spur further investigation. The scale and resolution of our atlas also enables the construction of a refined model of cell states and regulators in partially overlapped and paralleled primary and secondary myogenesis in the limb marked by different panels of regulators, with the identification and validation of MSC as a key player in muscle stem cell maintenance.

Our atlas also leverages spatial data by placing subtly distinct single cell clusters into their anatomical context, shedding light on their true identity. In particular, three clusters of cells with subtly different transcriptomes mapped to the distal limb with different distributions, which we term “distal” (LHX2^+^ MSX1^+^ SP9^+^), “RDH10^+^distal” (RDH10^+^ LHX2^+^ MSX1^+^) and “transitional” mesenchyme” (IRX1^+^ MSX1^+^). Similarly, two clusters in the tendon lineage map to the tendon and perimysium, giving insight into the subtle differences between these related tissues. Furthermore, through spatial transcriptomic analysis of the developing autopod, we connected physiological gene expression patterns to single gene health conditions that involve altered digital phenotype, demonstrating the clinical relevance of developmental cell atlas projects. We further maximised the utility of this study by presenting an integrated cross-species atlas with unified annotations as a resource for the developmental biology community that we expect will strengthen future studies of limb development and disease that utilise murine models.

One limitation in our study is the lack of samples from the earliest stages of limb bud development. Although we were able to dissect and process limbs from PCW5.1 onwards, the logistical limitations of working with human embryonic tissue precluded analysis of the nascent limb bud. This in turn prevented analysis of the earliest patterning and maturation events, such as the first wave of HOX gene expression. It also added to the difficulty of an unbiased cross-species comparison where more early-stage mouse samples are more accessible, let alone the data integration is already challenged by differences in lab protocols, transcriptome reference completeness and ortholog definitions etc. Furthermore, the small number of FGF8^+^AER cells captured by our experiments, together with the limited expression observed using RNA-ISH, suggest that during our sampling window, FGF8 expression in the distal ectoderm was already downregulated. In addition, whilst we were able to investigate FGFR2 expression patterns in the limb, it should be noted that the assays used are unable at present to distinguish between its IIIb (highest FGF10 affinity) and IIIc (highest FGF8 affinity) isoforms.

Whilst the combination of single cell and spatial transcriptomics is an established method for tissue atlasing, we recognise the challenges of combining different technologies. For example, our single-cell data captured large numbers of chondrocytes, including prehypertrophic chondrocytes which mapped to the mid-diaphysis of the forming bones. Interestingly, analysis of spatial gene expression revealed abundant collagen-X expression in these regions; the marker gene for mature, hypertrophic chondrocytes. Our scRNAseq experiments captured only n=14 cells expressing collagen-X; a disproportionately small number given the tissue area of expression on the visium. One possible explanation of this is that the method of permeabilisation and RNA capture with visium in this case was superior to the single cell tissue dissociation and droplet-based capture method for profiling matrix-rich tissues such as mature cartilage. Another important factor to consider is the breadth of cell capture with each technique. For scRNAseq, tissue samples were dissociated in order to produce single cell solutions. A typical solution in such an experiment contains hundreds of thousands of cells, only a fraction of which are loaded onto the chromium system, meaning rarer cell populations are unlikely to be sequenced. By contrast, visium captures RNA from all cells within each voxel across an entire tissue section, thus rare cell populations should still contribute to the overall RNA signal obtained from a voxel, which may in part explain why transcripts that were rare in the single cell data, such as COL10A1, are more abundant in the visium data. These factors may also explain the low numbers of SCX^+^/SOX9^+^ enthesis progenitor cells captured in our study. We expect these technical considerations to feed forward into future atlasing endeavours involving cartilage, bone and other dense tissues, or any tissue where rare cell types exist and an in-depth transcriptomic profile of such cells is desired.

Although the use of spatial transcriptomics in this atlas gives valuable anatomical context to sequenced single cells, at present this technique is still limited in its utility due to its 50*μ*m resolution, as well as significant dead space between voxels that are not sequenced. In the case of the fetal limb, this translates to up to 25 cells per voxel in histologically dense tissues such as cartilage, and finer structures such as the endothelium (Extended Data Fig. 6g,h) that showed an uncertain pattern of co-localization to be validated at higher resolution. Thus whilst deconvolution techniques such as those utilised in this study can allow a broad appreciation of cell type location, a true understanding of tissue architecture at the single cell, whole transcriptome level remains elusive. Furthermore, such a large sampling area makes fine-grain analyses of gene expression based on histology or anatomy challenging. In this study, we attempted to identify genes that vary between individual digits, but found very few. This is likely due to a lack of statistical power, with each digit only occupying 10-16 voxels.

## Methods

### Human tissue sample collection

First trimester human embryonic tissue was collected from elective termination of pregnancy procedures at Addenbrookes Hospital, Cambridge, UK under full ethical approval (REC-96/085; for scRNA-seq and Visium), at Guangzhou Women and Children’s Medical Center, China under the licence of ZSSOM-2019-075 and 2022-050A01 approved by the human research ethics committee of Sun Yat-sen University and Guangzhou Women and Children’s Medical Centre (for In-situ hybridisation and immunohistochemistry). For light-sheet fluorescence microscopy, tissues were obtained through INSERM’s HuDeCA Biobank and made available in accordance with the French bylaw. Permission to use human tissues was obtained from the French agency for biomedical research (Agence de la Biomédecine, Saint-Denis La Plaine, France; N° PFS19-012) and INSERM Ethics Committee (IRB00003888). Written, informed consent was given for tissue collection by all patients. Embryonic age (post conception weeks, PCW) was estimated using the independent measurement of the crown rump length (CRL), using the formula PCW (days) = 0.9022 × CRL (mm) + 27.372.

### Human tissue processing and scRNA-seq data generation

Embryonic limbs were dissected from the trunk under a microscope using sterile microsurgical instruments. In order to capture cells of the interzone, four samples (a hindlimb and a forelimb from both PCW5.6 and 6.1) were then further dissected into proximal, middle (containing undisturbed interzone) and distal thirds prior to dissociation. For the PCW5.1 sample, no further dissection was performed due to small size and the limb was dissociated as a whole. For all other samples, the limb was dissected into proximal and distal halves prior to dissociation.

Dissected tissues were mechanically chopped into a mash, and then were digested in Liberase TH solution (Roche, 05401135001, 50 *μ*g/ml) at 37°C for 30-40 min till no tissue pierce visible. Digested tissues were filtered through 40 *μ*m cell strainers followed by centrifugation at 750g for 5 min at 4°C. Cell pellets were resuspended with 2% FBS in PBS if the embryos were younger than PCW8, otherwise red blood cell lysis (eBioscience, 00-4300) was performed. The single-cell suspensions derived from each sample were then loaded onto separate channels of a Chromium 10x Genomics single cell 3’version 2 library chip as per the manufacturer’s protocol (10x Genomics; PN-120233). cDNA sequencing libraries were prepared as per the manufacturer’s protocol and sequenced using an Illumina Hi-seq 4000 with 2x150bp paired-end reads.

### Mouse tissue sample collection and scRNA-seq data generation

Timed pregnant C57BL/6J wild type mice were ordered from Jackson Laboratories. Embryos were collected at E12.5, E13.5 and E16.5. Only right side forelimbs and hindlimbs were used in this study: n=5 at the E12.5 timepoint, n=5 at E13.5 and n=2 at E16.5. Hindlimbs and forelimbs were pooled separately in ice cold HBSS (Gibco, 14175-095), and dissected into proximal, mid and distal limb regions, which were again separately pooled in 200 *μ*l of HBSS placed in a drop in the centre of a 6 cm culture plate. Tissues were then minced with a razor blade, and incubated with an addition of 120 *μ*l of diluted DNAse solution (Roche, 04716728001) at 37C for 15 minutes. DNAse solution: 1 ml UltraPure water (Invitrogen, 10977-015) 110 *μ*l 10X DNAse buffer, and 70 *μ*l DNAse stock solution. 2 ml of diluted Liberase TH (Roche, 05401151001) was then added to the plate, and the minced tissue suspension was pipetted into a 15 ml conical centrifuge tube. The culture plate was rinsed with 2 ml, and again with 1 ml of fresh Liberase TH which was serially collected and added to the cell suspension. The suspension was incubated at 37°C for 15 minutes, triturated with a P1000 tip, and incubated for an additional 15 minutes at 37°C. Liberase TH solution: 50X stock was prepared by adding 2 ml PBS to 5 mg of Liberase TH. Working solution is made by adding 100 *μ*l 50X stock to 4.9 ml PBS. After a final gentle trituration of the tissue with a P1000 tip, the suspension was spun at 380g in a swinging bucket rotor at 4°C for 5 minutes. After removing the supernatant, cells were resuspended in 5 ml of 2% fetal bovine serum in PBS, and filtered through a pre-wetted 40 *μ*m filter (Falcon, 352340). After spinning again at 380g at 4°C for 5 minutes, the supernatant was removed and cells were resuspended in 200 ul 2% FBS in PBS. A small aliquot was diluted 1:10 in 2% FBS/PBS and mixed with an equal volume of Trypan Blue for counting on a hemocytometer. The full suspension was diluted to 1.2 million cells/ml for processing on the 10x Genomics Chromium Controller, with a target of 8000 cells/library. Libraries were processed according to the manufacturer’s protocol, using the v3 Chromium reagents.

### Visium spatial transcriptomic experiments of human tissue

Whole embryonic limb samples at PCW6-8 were embedded in OCT within cryo wells and flash-frozen using an isopentane & dry ice slurry. Ten-micron thick cryosections were then cut in the desired plane and transferred onto Visium slides prior to haematoxylin and eosin staining and imaged at 20X magnification on a Hamamatsu Nanozoomer 2.0 HT Brightfield. These slides were then further processed according to the 10X Genomics Visium protocol, using a permeabilisation time of 18min for the PCW6 samples and 24 minutes for older samples. Images were exported as tiled tiffs for analysis. Dual-indexed libraries were prepared as in the 10X Genomics protocol, pooled at 2.25 nM and sequenced 4 samples per Illumina Novaseq SP flow cell with read lengths 28bp R1, 10bp i7 index, 10bp i5 index, 90bp R2.

### Digit region analysis of Visium data

The Visium data were clustered by Louvain algorithm after filtering genes that were expressed in less than 1 spot and performing normalisation and logarithmisation. After that, the spot clusters of interest were annotated based on H&E histology and marker genes. The differential expression testing was performed by Wilcoxon test using Scanpy (sc.tl.rank_gene_group).

### Alignment, quantification and quality control of scRNA-seq data

Droplet-based (10X) sequencing data were aligned and quantified using the Cell Ranger Single-Cell Software Suite (v3.0.2, 10X Genomics). The human reference is the hg38 genome refdata-cellranger-GRCh38-3.0.0, available at: http://cf.10xgenomics.com/supp/cell-exp/refdata-cellranger-GRCh38-3.0.0.tar.gz. The mouse reference is the mm10 reference genome refdata-gex-mm10-2020-A, available at: https://cf.10xgenomics.com/supp/cell-exp/refdata-gex-mm10-2020-A.tar.gz. The following quality control steps were performed: (i) cells that expressed fewer than 200 genes (low quality) were excluded; (ii) genes expressed by less than 5 cells were removed; (iii) cells in which over 10% of unique molecular identifier (UMIs) were derived from the mitochondrial genome were removed.

### Alignment and quantification of human Visium data

Raw FASTQ files and histology images were processed, aligned and quantified by sample using the Space Ranger software v.1.2.0. which uses STAR v.2.5.1b52 for genome alignment, against the Cell Ranger hg38 reference genome refdata-cellranger-GRCh38-3.0.0, available at: http://cf.10xgenomics.com/supp/cell-exp/refdata-cellranger-GRCh38-3.0.0.tar.gz.

### Doublet detection of scRNA-seq data

Doublets were detected with an approach adapted from a previous study^159^. In the first step of the process, each 10X lane was processed independently using the Scrublet to obtain per-cell doublet scores. In the second step of the process, the standard Scanpy processing pipeline was performed up to the clustering stage, using default parameters^160^. Each cluster was subsequently separately clustered again, yielding an over-clustered manifold, and each of the resulting clusters had its Scrublet scores replaced by the median of the observed values. The resulting scores were assessed for statistical significance, with P values computed using a right tailed test from a normal distribution centred on the score median and a median absolute deviation (MAD)-derived standard deviation estimate. The MAD was computed from above-median values to circumvent zero truncation. The P values were corrected for false discovery rate with the Benjamini–Hochberg procedure and were used to assess doublet level. The clusters from batch-corrected overall clustering across all the samples that have median score lower than 0.1 and are supported by absence of exclusive marker genes or literature are manually curated and removed (1,450 doublets were removed in human data, 958 in murine data).

### Data preprocessing and integration of scRNA-seq data

Preprocessing included data normalisation (pp.normalize_per_cell with 10,000 counts per cell after normalization), logarithmise (pp.log1p), highly variable genes (HVGs) detection (pp.highly_variable_genes and select for highly correlated ones as described in ^152^) per batch and merging, data feature scaling (pp.scale), cell cycle and technical variance regressing (tl.score_gene_cell_cycle and pp.regress_out(adata,[’S_score’, ’G2M_score’,’n_counts’,’percent_mito’])), and principal component analysis (PCA) (tl.pca with 100 components) performed using the Python package Scanpy (v.1.8.2). bbknn (v.1.5.1) was used to correct for batch effect between sample identities with the following parameters (n_pcs = 100, metric= “euclidean”, neighbors_within_batch=3, trim=299, approx=False). Following this, further dimension reduction was performed using Uniform Manifold Approximation and Projection (UMAP) (scanpy tl.umap with default parameters) based on the corrected neighbohood graph of bbknn.

### Clustering and annotation of scRNA-seq data

We first applied Leiden graph-based clustering (scanpy tl.leiden with default parameters) to perform unsupervised cell classification. Each cluster was then subclustered if heterogeneity is still observed and was manually annotated (Extended Data Table 1 for marker genes) and curated as described before ^16^.To make sure all the curated Leiden clusters could clearly be mapped onto their UMAP embedding coordinates, we performed the partition-based graph abstraction (PAGA) (tl.paga with the Leiden clusters) and rerun UMAP with the initial position from PAGA.

### Deconvolution of human Visium data-cell2location

To map clusters of cells identified by scRNA-seq in the profiled spatial transcriptomics slides, we used the cell2location method ^20^. In brief, this involved first training a negative binomial regression model to estimate reference transcriptomic profiles for all the scRNA-seq clusters in the developing limb. Next, lowly expressed genes are excluded as per recommendations for use of cell2location, leaving 13,763 genes for downstream analysis. Next, we estimated the abundance of each cluster in the spatial transcriptomics slides using the reference transcriptomic profiles of different clusters. This was applied to all slides simultaneously, using the sample id as the batch_key and categorical_covariate_keys. To identify microenvironments of co-localising cell clusters, we used non-negative matrix factorisation (NMF) implementation in scikit-learn, utilising the wrapper in the cell2location package ^161^. A cell type was considered part of a microenvironment if the fraction of that cell type in said environment was over 0.2.

### Alignment and merging of multiple visium sections

In order to analyse the whole PCW 8.1 human hindlimb, we took three consecutive ten micron sections from different regions and placed them on different capture areas of the same Visium LP slide. The first section spanned the distal femur, knee joint and proximal tibia (sample C42A1), the second the proximal thigh (C42B1) and the third the distal tibia, ankle and foot (C42C1).

The images from these 3 visium capture areas were then aligned using the TrackEM plugin (Fiji)^162^. Following affine transformations of C42B1 and C42C1 to C42A1, the transformation matrices were exported to an in-house pipeline (/22-03-29-visium-stitch/limb_reconst.ipynb) for complementary alignment of the spot positions from the SpaceRanger output to the reconstructed space. In addition we arbitrarily decided that for regions of overlapping spots we kept the spots from the centre portion (see supp) and the same decision was made for the reconstructed image in order to maintain the 1-1 relationship between the image and genetic profile. Next we merged the 3 library files by an in-house pipeline (22-03-29-visium-stitch/N1-join-limb.ipynb) and matched the reconstructed image to the uniform AnnData object.

### Trajectory analysis of human scRNA-seq data

Development trajectories were inferred by combining diffusion maps (DM), PAGA and force-directed graph (FDG). The first step of this process was to perform the first nonlinear dimensionality reduction using DM (scanpy tl.diffmap with 15 components) and recompute the neighborhood graph (scanpy pp.neighbors) based on the 15 components of DM. In the second step of this process, PAGA (scanpy tl.paga) was performed to generate an abstracted graph of partitions. Finally, FDG was performed with the initial position from PAGA (scanpt tl.draw_graph) to visualise the development trajectories.

### RNA Velocity calculations for mesenchymal compartment

The scVelo version 0.24 package for Python was used to calculate a ratio of spliced: unspliced mRNA abundances in the dataset ^163^. The data was sub-clustered to the mesenchymal compartment for a single sample (PCW7.2). The data was then processed using default parameters following preprocessing as described in Scanpy scVelo implementation. The samples were pre-processed using functions for detection of minimum number of counts, filtering and normalisation using scv.pp.filter_and_normalise and followed by scv.pp.moments function using default parameters. The gene specific velocities were then calculated using scv.tl.velocity with mode set to stochastic, and scv.tl.velocity_graph functions and visualised using scv.pl.velocity_graph function.

### Cell-cell communication analysis of human scRNA-seq data

Cell–cell communication analysis was performed using the CellPhoneDB (v.2.1.4) for each dataset at the same stage of development. The stage matched Visium data was used to validate the spatial distance and expression pattern of significant (P < 0.05) ligand-receptor interactions.

### Regulon analysis of transcription factors (TFs)

To carry out transcription factor network inference, Analysis was performed as previously described^164^ using the pySCENIC python package (v.0.10.3). For the input data, we filtered out the genes that were expressed in less than 10 percent of the cells in each cell cluster. Then, we performed the standard procedure including deriving co-expression modules (pyscenic grn), finding enriched motifs (pyscenic ctx) and quantifying activity (pyscenic aucell).

### Integration of human and mouse scRNA-seq data

Mouse orthologs were first “translated” to human genes using MGI homology database (https://www.informatics.jax.org/homology.shtml). Processed human and mouse data were then merged together using outer join of all the genes The matched dataset was then integrated by MultiMAP using the MultiMAP_Integration() function, using separately pre-calculated PCs and the union set of previously calculated mouse and human feature genes (including both orthologs and non-orthologs) to maximise biological variance. Downstream clustering and embedding were performed as usual and cell-type annotation was based on marker genes. Cell-type composition of proximal, middle and distal segments of the same limb was visualised using plotly.express.scatter_ternary() function. To capture the differential expression of sparsely captured genes, the odds ratio of the percentages of non-zero cells between groups of cells was used to select for proximal/distal or fore/hind biassed genes with a cutoff at 30 fold and 3 fold respectively.

### Immunohistochemistry

The lower limbs were post-fixed in 4% PFA for 24 h at 4°C followed by paraffin embedding. A thickness of 4 *μ*m sections were boiled in 0.01 M citrate buffer (pH 6.0) after dewaxing. Immunofluorescence staining was then carried out as described previously^165^. Primary antibodies for RUNX2 (1:50, Santa Cruz, sc-390715), THBS2 (1:100, Thermo Fisher, PA5-76418), COL2A1 (1:200, Santa Cruz, sc-52658), PITX1 (1:30, Abcam, Ab244308), PAX3 (1:1, DSHB, AB_528426 supernatant) and ALDH1A3 (1:50, Proteintech, 25167-1-AP) and MYH3 (1:3, DSHB, AB_528358 supernatant) were incubated overnight at 4°C. After washing, sections were incubated with appropriate secondary antibodies Alexa Flour 488 goat anti-mouse IgG1 (Invitrogen, A-21121), Alexa Flour 647 goat anti-mouse IgG2b (Invitrogen, A-21242), Alexa Flour 488 goat anti-mouse IgG (H+L) (Invitrogen, A-11029) and Alexa Flour 546 goat anti-rabbit IgG (H+L) (Invitrogen, A-11035) at room temperature for 1h, and were mounted using FluorSave Reagent (Calbiochem, 345789). For 3, 3 - diaminobenzidine (DAB) staining, we used Streptavidin-Peroxidase broad spectrum kit (Bioss, SP-0022) and DAB solution (ZSGB-BIO, ZLI-9017) following the manufacturers’ manuals. Primary antibodies PI16 (1:500, Sigma-Aldrich, HPA043763), FGF19 (1:500, Affinity, DF2651) and NEFH (1:1000, Cell Signaling, 2836) were applied. Single-plane images were acquired using an inverted microscope (Leica, DMi8).

### RNA in-situ hybridization

Fresh tissue samples were embedded in OCT and frozen at −80°C until analysis. Cryosections were cut at a thickness of 10 *μ*m or 12*μ*m using a cryostat (Leica CM1950 or CM3050). Prior to staining, tissue sections were post-fixed in 4% PFA for 15 min at 4°C. After a series of 50%, 70%, 100%, and 100% ethanol dehydration for 5 min each, tissue sections were treated with hydrogen peroxide for 10 min. Next, the sections were digested with protease IV (ACD, 322336) for 20-30 min at room temperature, alternatively, were digested with protease III (ACD, 322337) for 15 min after heat-induced epitope retrieval. RNA-ISH was then carried out manually or using BOND RX (Leica) by employing RNAscope® Multiplex Fluorescent Reagent Kit v2 Assay (ACD, 323110) or PinpoRNA^TM^ multiplex fluorescent RNA in-situ hybridization kit (GD Pinpoease, PIF2000) according to the manufacturers’ instructions. To visualize targeted RNAs from individual channels, different tyramide signal amplification (TSA) fluorescent substrates were incubated. Two sets of fluorophores TSA 520, TSA 570, TSA 650 (PANOVUE) and Opal 520, Opal 570, Opal 650 (Akoya Biosciences) were used and consistent results were obtained. For the staining of 4 probes, RNAscope® 4-plex Ancillary Kit (ACD, 323120) was applied additionally, and a combination of fluorophores TSA520, TSA570, Opal620, and Opal690 were used. The stained sections were imaged with either AxioScan.Z1 (ZEISS) or Opera® Phenix™ High-Content Screening System (PerkinElmer).

### Light Sheet Fluorescence Microscopy

Embryonic and fetal limbs were dissected from morphologically normal specimens collected from PCW5 to 6.5. Candidate antibodies were screened by immunofluorescence on cryosections obtained from OCT-embedded specimens as previously described^166^. Whole-mount immunostaining of the limbs was performed as previously described, with primary antibody incubation at 37°C reduced to 3 days followed by 1 days in secondary antibodies. Samples were (embedded in 1.5% agarose) and optically cleared with solvents using the iDisco+ method. Cleared samples were imaged with a Blaze light-sheet microscope (Miltenyi Biotec) equipped with sCMOS camera 5.5MP controlled by Imspector Pro 7.5.3 acquisition software. A 12x objective with 0.6X or 1X magnification (MI Plan NA 0.53) was used. Imaris (v10.0, BitPlane) was used for image conversion, processing, and video production. All raw image data are available on request (AC, RB).

### List of the Antibodies for LSFM

IRX1 Sigma-Aldrich Cat# HPA043160, RRID:AB_10794771 (1/200e)

MSX1 R&D Systems Cat# AF5045, RRID:AB_2148804 (1/500e)

LHX2 Abcam Cat# ab184337 (1/1000e)

SOX9 Abcam Cat# ab196184, RRID:AB_2813853 (1/500e)

MAFB Abcam Cat# ab223744, RRID:AB_2894837 (1/500e)

Donkey Anti-Rabbit IgG H&L (Alexa Fluor® 555) Abcam Cat# ab150062, RRID:AB_2801638 (1/800e)

Donkey Anti-Goat IgG H&L (Alexa Fluor® 790) Abcam Cat# ab175745 (1/300e)

### MSC knockdown in human primary myoblasts

#### Isolation of human primary myoblast cells

The thighs from human embryos were processed as described^167^, except that the dissociated cells were not treated with erythrocyte lysis solution, and were incubated with anti-human CD31 (eBioscience, 12-0319-41), CD45 (eBioscience, 12-0459-41) and CD184 (eBioscience, 17-9999-41) antibodies for cell sorting. Fluorescent activated cell sorting (FACS, BD, influx) sorted CD31-CD45-CD184+ cells were cultured in complete growth medium DMEM supplemented with 20% FCS and 1% penicillin/streptomycin (Gibco, 15140122).

#### siRNA transfection

Human primary myoblasts were seeded into a 6-well plate one night before transfection. When the cell density reached approximately 50% confluence, oligos of small interfering RNA (siRNA) against MSC (si-MSC) and negative control (NC) were transfected using Lipofectamine 3000 reagent (Invitrogen, L3000015) at a final concentration of 37.5 nM. After incubation for 16 h, the growth medium was replaced with differentiation medium containing 2% horse serum and 1% penicillin/streptomycin in DMEM. After culturing for an additional 6-8 h, the cells were collected for RNA extraction. Initially three siRNA oligos (Bioneer, 9242-1, 2, 3) were tested, and the third one with sense sequences 5*′*-GAAGUUUCCGCAGCCAACA-3*′* were used in this study.

#### RNA extraction and quantitative PCR (qPCR)

Total cell RNA was extracted with the EZ-press RNA purification kit (EZBioscience, B0004D), and the cDNA was synthetised using the PrimeScript RT Master Mix Kit (TaKaRa, RR036A). The qPCR was performed using PerfectStartTM Green qPCR Super Mix (TransGen Biotech, 1) on a Real-time PCR Detection System (Roche, LightCycle480 II). RPLP0 served as an internal control, and the fold enrichment was calculated using the formula 2*−ΔΔCt*. The following primers (5’-3’) were used:

RPLP0 forward: ATGCAGCAGATCCGCATGT, reverse:

TTGCGCATCATGGTGTTCTT;

MSC forward: CAGGAGGACCGCTATGAGAA, reverse:

GCGGTGGTTCCACATAGTCT;

MYOG forward: AGTGCCATCCAGTACATCGAGC, reverse:

AGGCGCTGTGAGAGCTGCATTC;

MYH2 forward: GGAGGACAAAGTCAACACCCTG, reverse:

GCCCTTTCTAGGTCCATGCGAA;

MYH3 forward: CTGGAGGATGAATGCTCAGAGC, reverse:

CCCAGAGAGTTCCTCAGTAAGG;

MYH4 forward: CGGGAGGTTCACACAAAAGTCATA, reverse:

CCTTGATATACAGGACAGTGACAA;

TNNT1 forward: AACGCGAACGTCAGGCTAAGCT, reverse:

CTTGACCAGGTAGCCGCCAAAA.

**Extended Figure 1.**
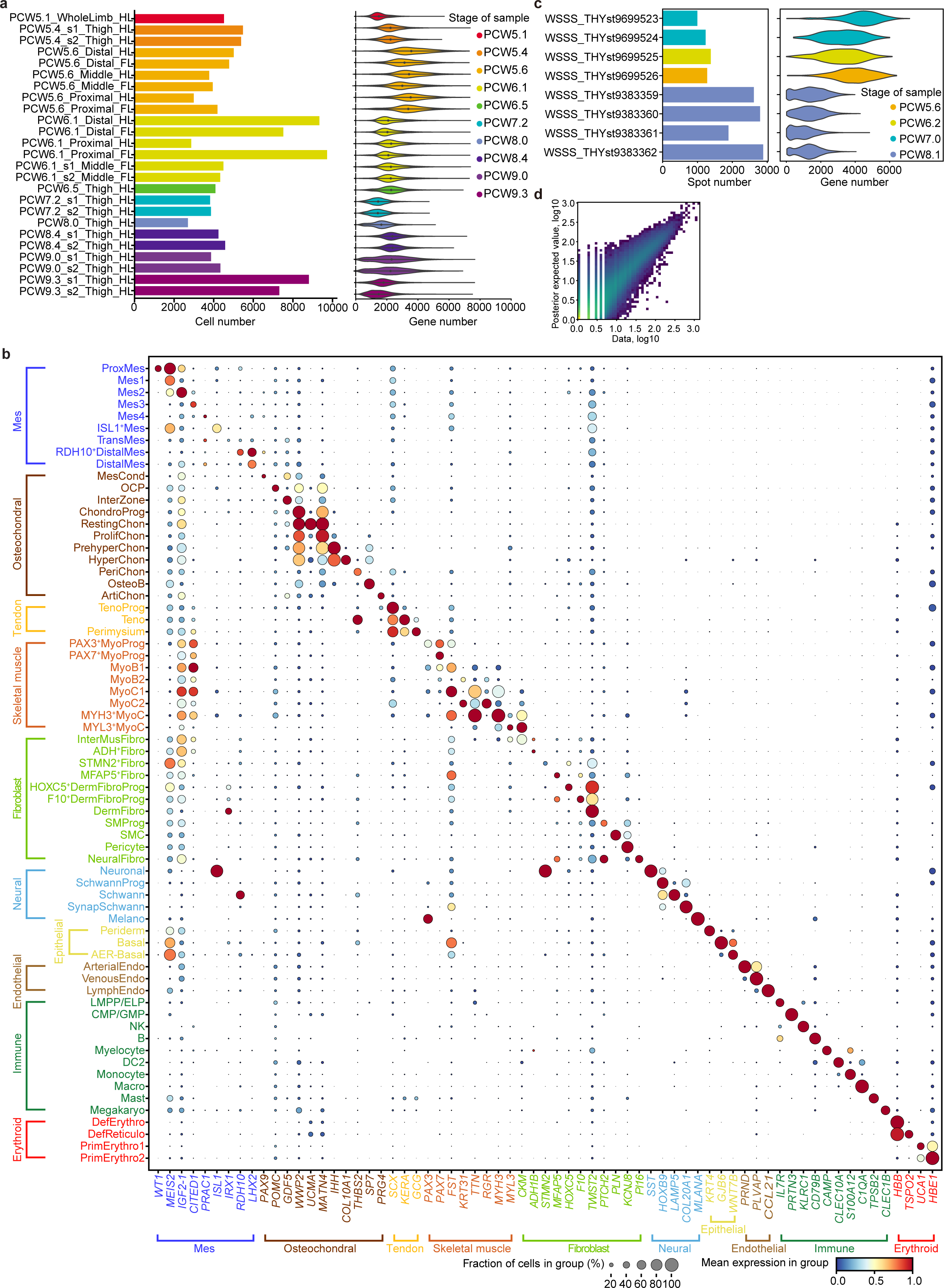
Data quality and preprocess of human scRNA-seq and spatial visium data. **a**, Bar plot and violin plot showing the sample size and per-cell quality of each library, separately, coloured by stage. **b**, Dot plot showing the expression level of marker genes for each cell cluster. The colour bar indicates the linearly scaled mean of expression level. **c**, Bar plot and violin plot showing the number of voxels and per-voxel quality of each 10X Visium library, separately, coloured by stage. **d**, Scatter plot showing the reconstruction accuracy of cell2location.

**Extended Figure 2.**
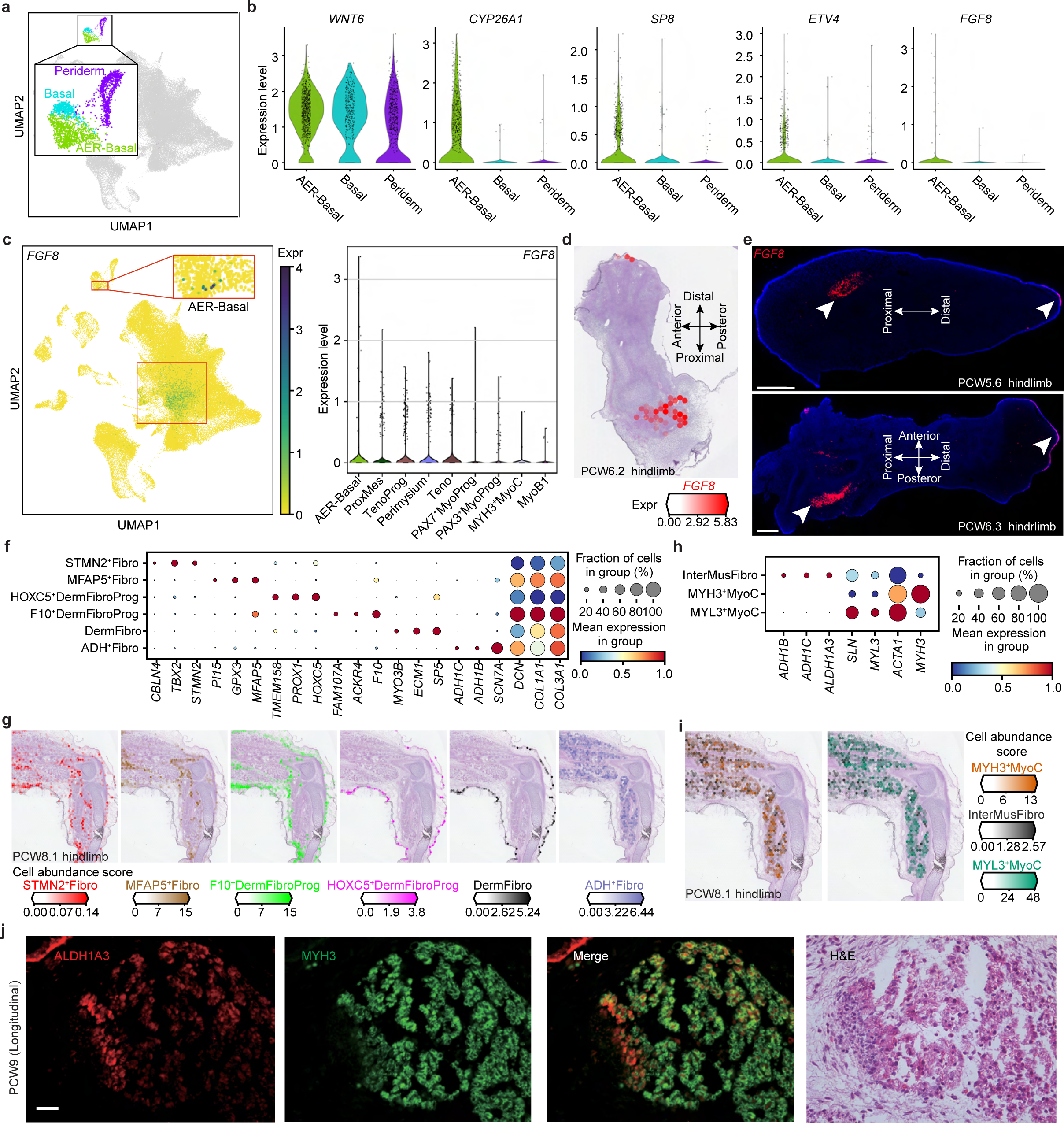
The heterogeneity of epidermis and fibroblast. **a**, Uniform manifold approximation and projection (UMAP) visualization of AER-basal, basal and periderm cells. **b**, Violin plot showing the expression level of WNT6, CYP26A1, SP8, ETV4 and FGF8 in AER-basal, basal and periderm cells. The expression value has been normalized and log-transformed. **c**, UMAP (left panel) and violin plot (right panel) showing the expression level of FGF8 in the human limb. ProxMes, proximal mesenchyme; TenoProg, tendon progenitor; MyoProg, myogenic progenitor; MyoB, myoblast; MyoC, Myocyte. **d**, Heatmap across tissue section from PCW6.2 (post conception week 6 plus 2 days) human hindlimb showing FGF8 expression. **e**, Fluorescence staining of tissue sections from human hind limb showing the expression pattern of FGF8 in situ. Scale bar, 500um. **f**, **h**, Dot plot showing the expression level of marker genes for selected fibroblast (Fibro) clusters (f) and muscle interstitial fibroblast (InterMusFibro) (h). The colour bar indicates the linearly scaled values of expression level. DermFibro, dermal Fibro; DermFibroProg, DermFibro progenitor. **g**, **i**, Heatmaps across tissue sections from PCW8.1 human hindlimb showing inferred abundance of each fibroblast cluster (g) and InterMusFibro (i). **j**, Immunofluorescence staining of MYH3 and ALDH1A3 on the skeletal muscle tissue (as also shown by H&E staining) from a PCW9 longitudinal section. Scale bar, 50 *μ*m.

**Extended Figure 3.**
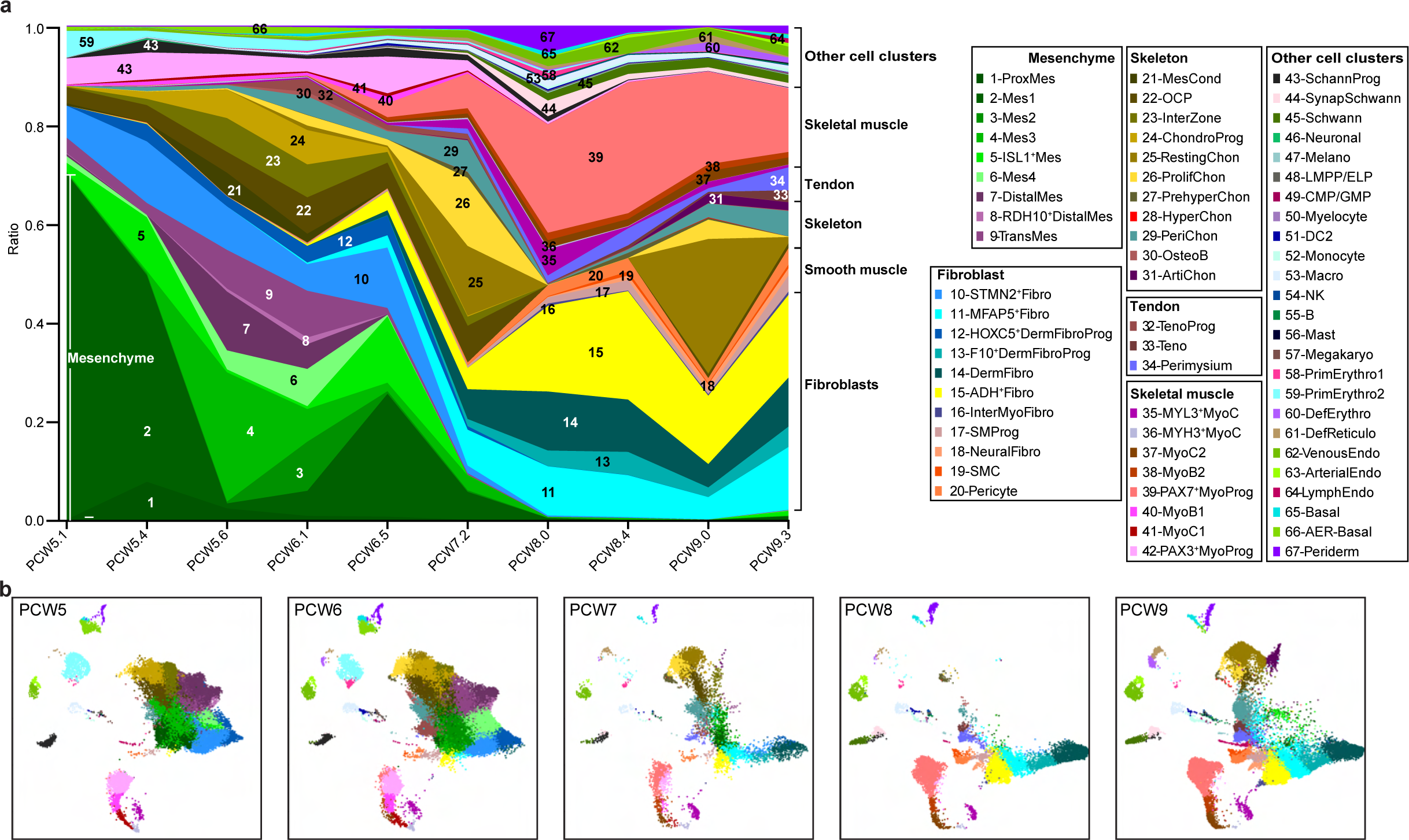
Dynamic changes in cell types of the human embryonic limb over developmental time. **a**, Fraction of cell type per time point, coloured by cell type and grouped by tissue type. Cluster abbreviations same as Fig. 1. **b**, Uniform manifold approximation and projection (UMAP) visualisation of cells per post conception week (PCW), coloured by cell type in a.

**Extended Figure 4.**
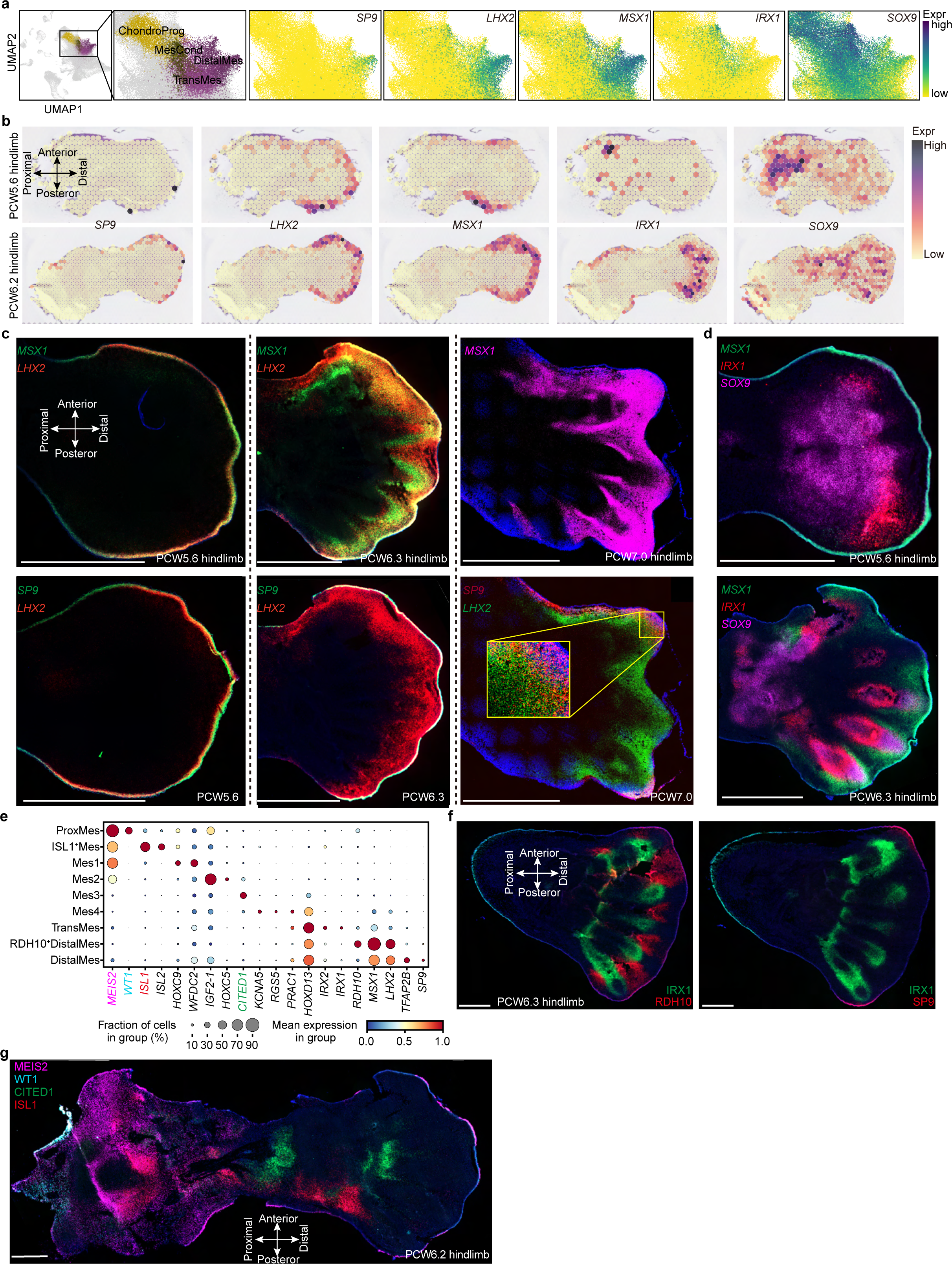
The heterogeneity of Mesenchyme. **a**, Uniform manifold approximation and projection (UMAP) plot showing the cell clusters of Chondrogenic progenitor (ChondroProg), mesenchymal condensate cell (MesCond), transitional mesenchyme (TransMes) and distal mesenchyme (DistalMes) (left panel) as well as the expression level of SP9, LHX2, MSX1, IRX1 and SOX9. The color bar indicates the normalized and log-transformed expression values. **b**, Heatmaps across tissue sections from human hindlimb at stage of PCW5.6 (post conception week 6 plus 2 days) and PCW6.2 showing the spatial expression pattern of SP9, LHX2, MSX1, IRX1 and SOX9. The color bar indicates the normalized log-transformed expression values. **c**, **d**, Fluorescence staining of tissue sections from human hind limb showing the spatial expression pattern of SP9, LHX2 and MSX1 (c), as well as MSX1, IRX1 and SOX9 (d) at different stage. Scale bar, 1mm. **e**, Dot plot showing the expression level of marker genes for different cell clusters of mesenchyme. The color bar indicates the linearly scaled mean of expression level. ProxMes, proximal Mes. **f**, **g**, RNA-ISH of tissue sections from human hindlimb showing the spatial expression pattern of *MEIS2, WT1, CITED1 and ISL1* (f), as well as *IRX1, SP9 and RDH10* (g) Scale bar, 1mm.

**Extended Figure 5.**
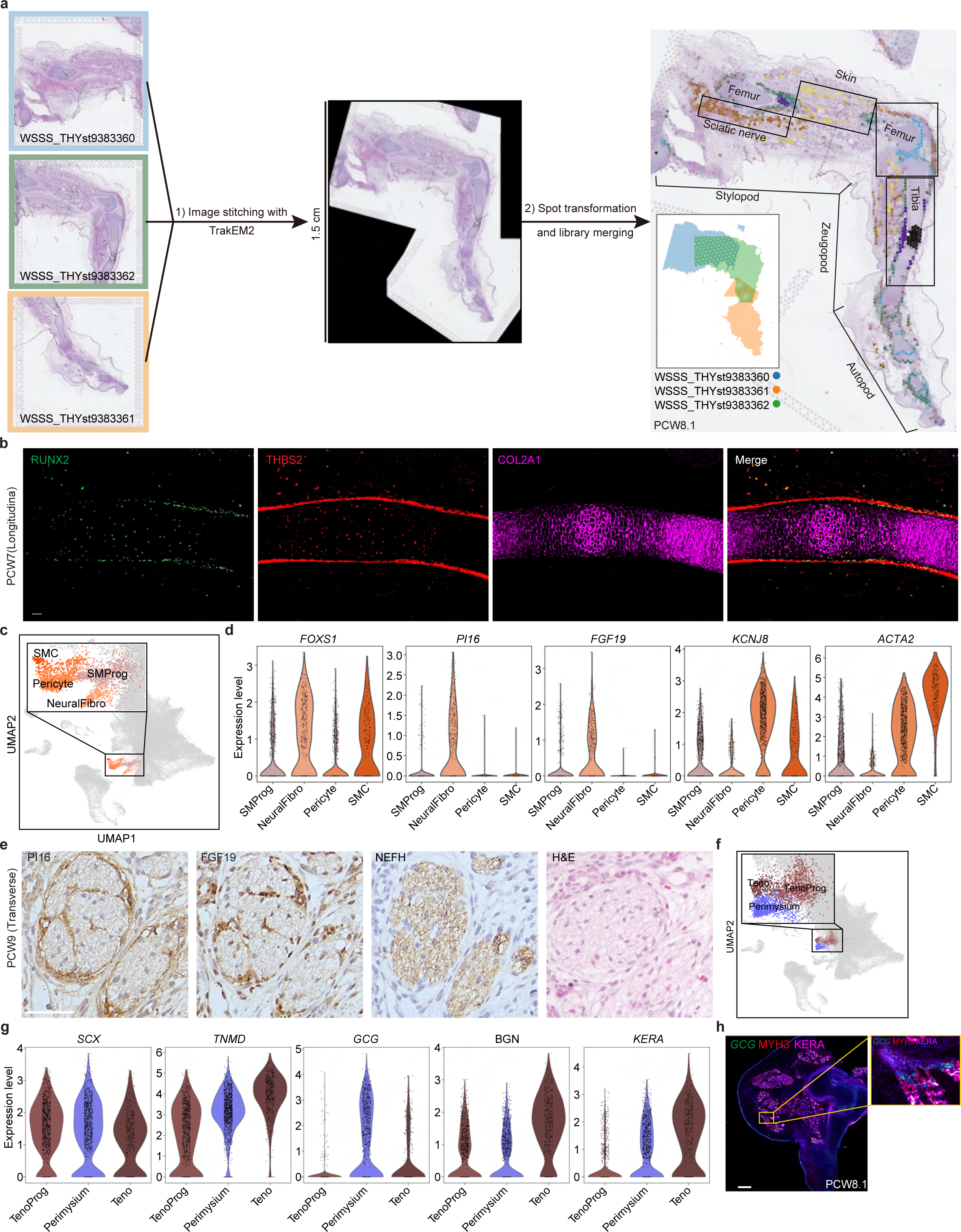
Identification of novel cell types at spatial and single-cell level. **a**, The workflow for merging the 10X Visium spatial data of the human limb. **b**, Immunofuorescence staining of RUNX2, THBS2 and COL2A1 on the longitudinal section of the tibia from a PCW7 embryo. Scale bar, 50 *μ*m. **c**, Uniform manifold approximation and projection (UMAP) visualization of smooth muscle progenitor (SMProg), neural fibroblasts (NeuralFibro), pericyte and smooth muscle (SMC). **d**, Violin plot showing the expression level of FOXS1, PI16, FGF19, KCNJ8 and ACTA2 in SMProg, NeuralFibro, pericyte and SMC, using normalised and log-transformed values. **e**, Immunohistochemical staining of PI16 and FGF19 showing the NeuralFibro in the sciatic nerve at PCW9. The neurofilament was stained with NEFH antibody. A neighbouring section stained with H&E solution is also shown. Scale bar, 50 *μ*m. **f**, UMAP visualization of tendon progenitor (TenoProg), tenocytes (Teno) and perimysium cells. **g**, Violin plot showing the expression level of SCX, TNMD, GCG, BGN and KERA in TenoProg, Perimysium, and Teno. The expression level of genes is the normalised and log-transformed values. **h**, RNA-ISH combined with immunohistochemistry of tissue sections from human hind limb showing the spatial expression pattern of GCG, MYH3 and KERA in situ. Scale bar, 1mm.

**Extended Figure 6.**
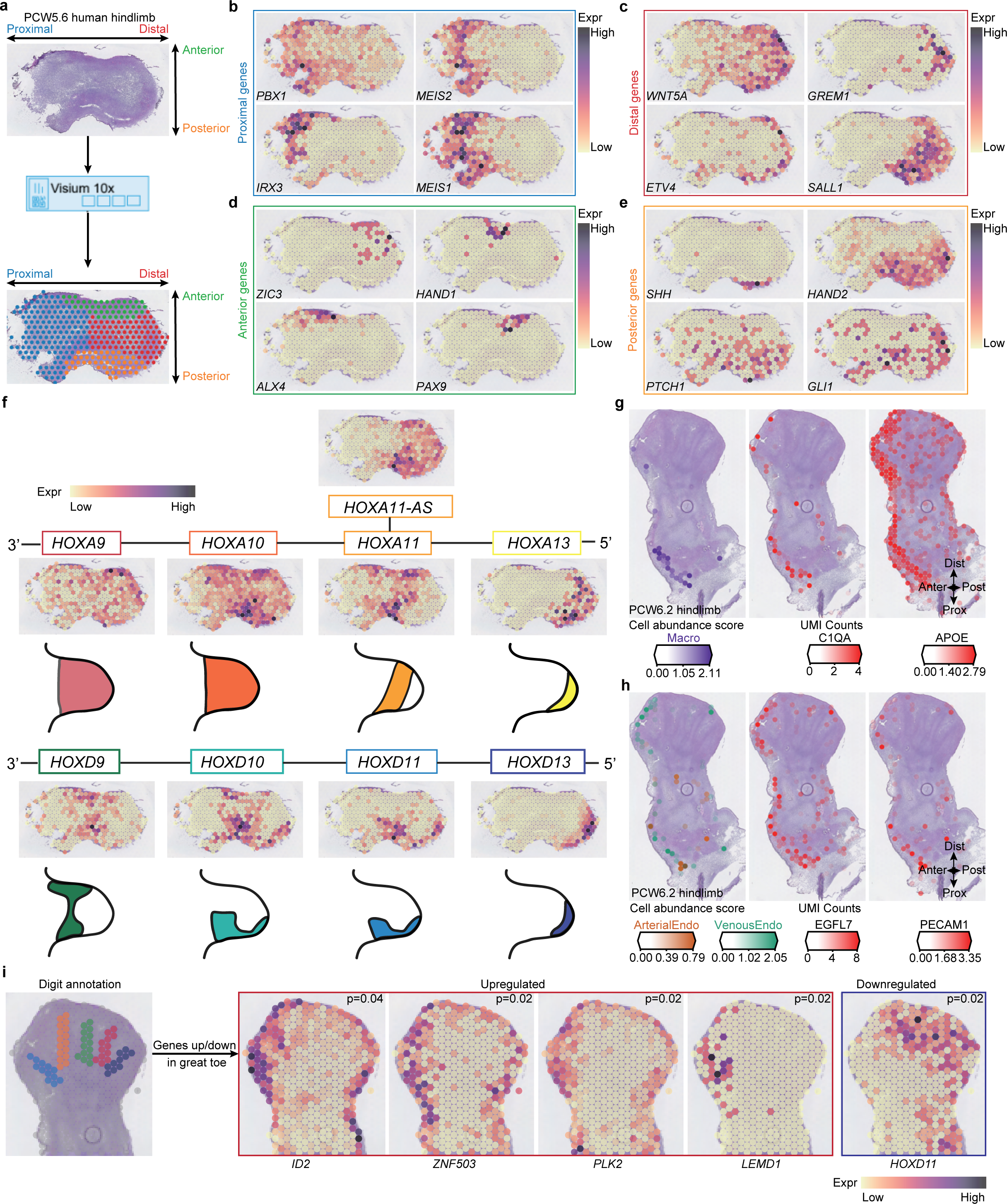
Spatial expression patterns of genes that determine human limb axis formation and morphogenesis. **a**, Overview of analysis workflow to identify genes specific to spatial location. **b-e**, Heatmaps across tissue section from the human hindlimb at stage of PCW5.6 showing spatial expression pattern of genes specific to proximal (b), distal (c), anterior (d) or posterior (e) regions. The expression level of genes is the normalised and log-transformed values. **f**, Heatmaps across tissue section from human hindlimb at stage of PCW5.6 showing spatial expression pattern of homeobox (HOX) A (top panel) and D (bottom panel) family genes. **g-h**, Heatmaps across tissue sections from the human hindlimb at stage PCW6.2 showing inferred abundance of macrophage (g) and endothelial cells (vein endothelial cells (VeinEndo) and arterial endothelial cells (ArterialEndo), h) as well as expression of maker genes. Anter, anterior; Post, posterior; Prox, proximal; Dist, distal. The expression level of genes is the normalized and logarithmic value of raw counts. **i**, Spatially resolved heatmaps across tissue section from the human hindlimb at stage of PCW5.6 showing spatial expression pattern of digit-associated genes in normalized and log-transformed values.

**Extended Figure 7.**
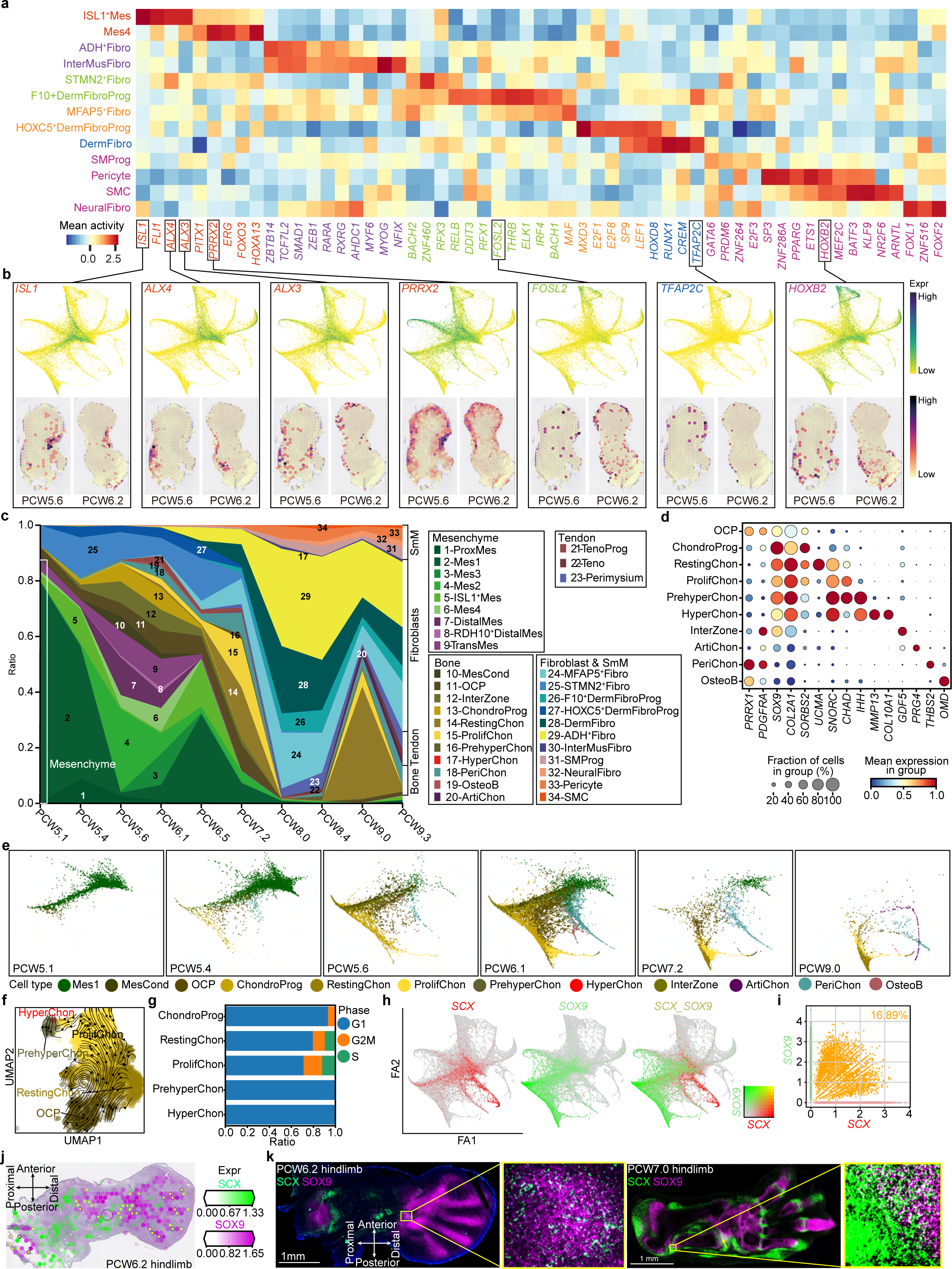
The transcriptional regulation of LPM differentiation in the human limb. **a**, Heatmap illustrating the vertically normalised mean activity of selected TFs for each cell type from soft connective lineage of LPM. **b**, Force-directed graphs (top panel) and heatmaps across tissue section from the human hindlimb at PCW5.6 and PCW6.2 (bottom panel) showing the expression pattern of representative TFs in normalised and log-transformed values. **c**, Stacked bar chart showing the fraction of cell type per time point, coloured by cell type and grouped by tissue type. PCW, post conception week. SmM, smooth muscle group; other abbreviations as per Figure 1; **d**, Dot plot showing the expression level of marker genes of osteochondral cell clusters. The colour bar indicates the mean normalised expression level. **e**, Force directed graph of cells per time point, coloured by cell type in a. **f**, Uniform manifold approximation and projection (UMAP) visualization of the chondrocyte lineage with arrows representing inferred differentiation directions (See Methods). **g**, Stacked bar chart showing the fraction of phase of cell cycle per osteochondral cell cluster. **h**, Dot plot showing the expression level of SCX and SOX9 in cell clusters. The colour bar indicates the linearly scaled mean of expression level. **i**, Heatmap across tissue section from the human hindlimb at PCW6.2 showing SCX and SOX9 expression in normalised and log-transformed values. The voxels marked with yellow asterisks express both SCX and SOX9. **j**, RNA-ISH of tissue sections from the human hindlimb showing co-expression of SCX and SOX9 in situ.

**Extended Figure 8.**
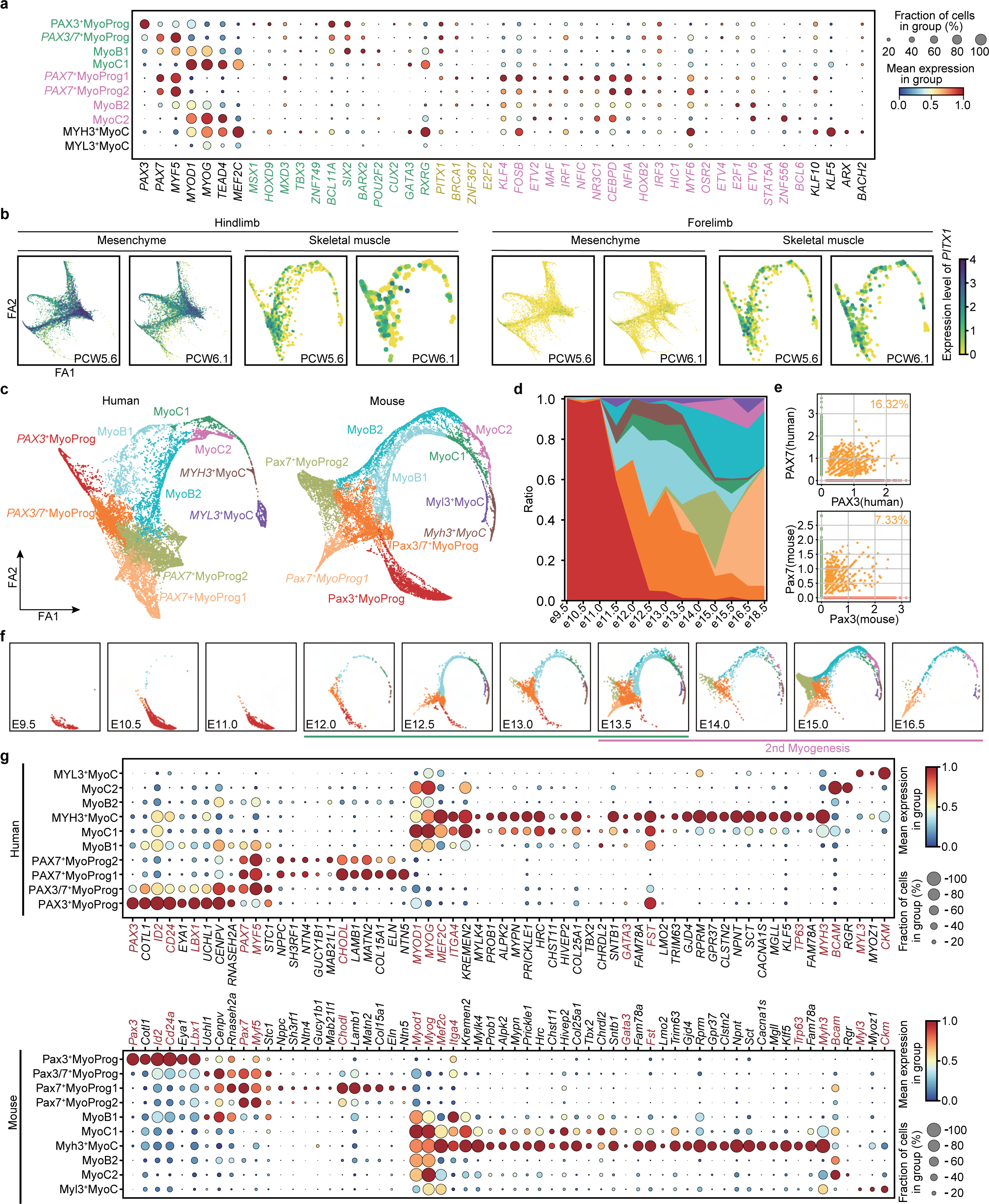
The transcriptional regulation of myogenesis in human and mouse. **a**, Dot plots showing expression level of TFs per cell cluster in humans, coloured by group (green, first myogenesis; pink, second myogenesis; yellow, both). The colour bar indicates the linearly scaled mean of expression level. MyoProg, myogenic progenitor; MyoB, myoblast; MyoC, myocyte. b *Force-directed graph showing the expression of PITX1 between forelimb (left panel) and hindlimb (right panel) in cells derived from mesenchyme and skeletal muscle lineage. The expression level of genes is the normalized and logarithmic value of raw counts.* **c**, Force-directed graph of human and mouse skeletal muscle cells, colored by cell clusters. **d**, Stacked bar chart showing the fraction of mouse cell clusters per time point, followed by the color code of mouse cell clusters in c. **e**, Scatter plots of PAX3 and PAX7 expression in mouse and human skeletal muscle cells. The percentages of double positive cells are given. **f**, Force-directed graph of mouse cells per time point, coloured by cell cluster in c. **g**, Dotplot of selected transcription factors expressed in human and mouse.

**Extended Figure 9.**
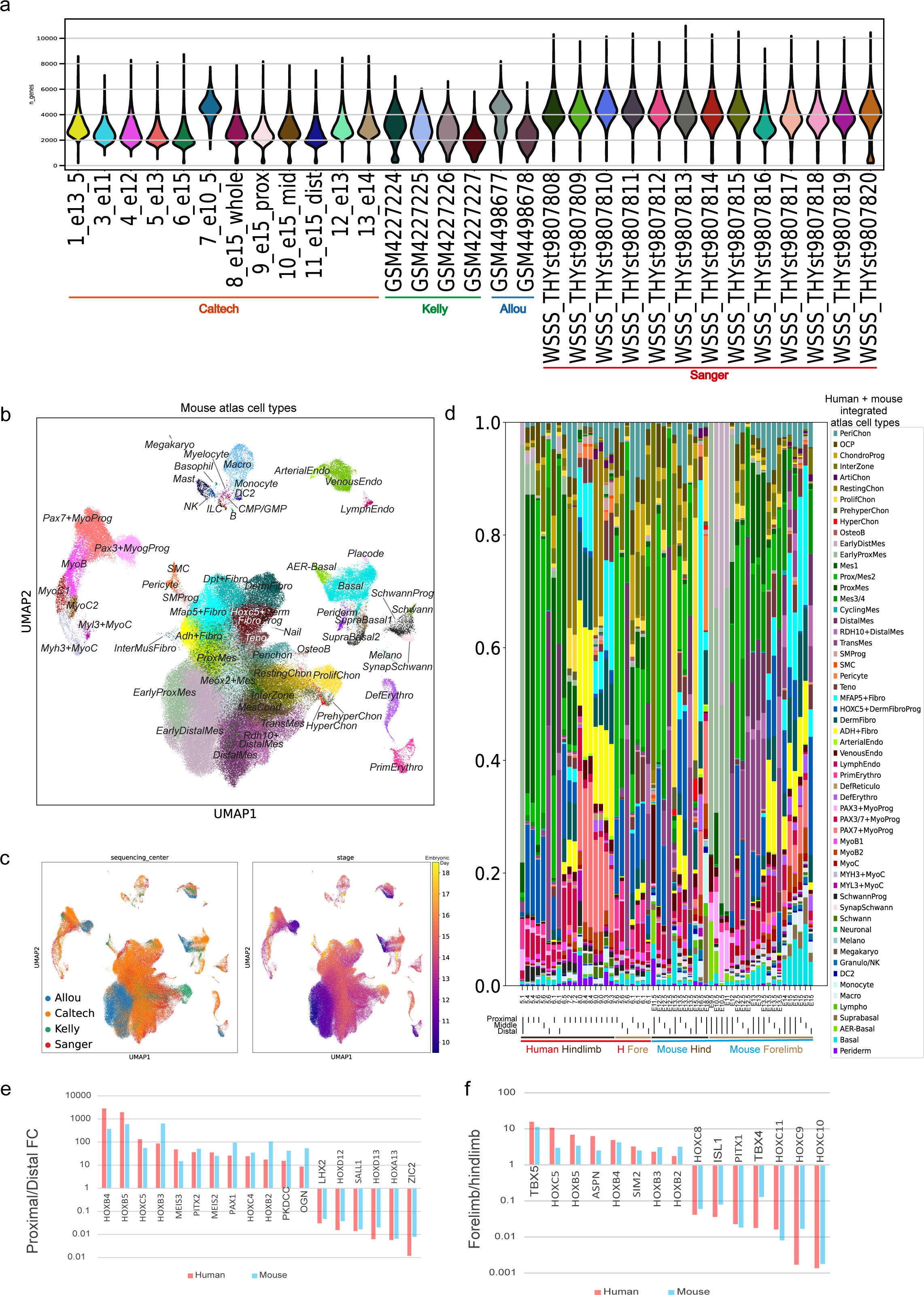
Comparing human and mouse embryonic limbs. **a**, Violin plots of sample quality for all the scRNA-seq data in the integrated atlas, coloured by library ID and group by sequencing centre at the bottom. **b-c**, The integrated mouse scRNA-seq data projected on a shared UMAP plane, coloured by cell clusters (b) or metadata (c). **d**, Cell-cluster proportions of each scRNA-seq library with dissection region, location and species labelled at the bottom. **e**, Genes enriched in proximal or distal segments in human and mouse. **f**, Genes enriched in the forelimb or hindlimb in human and mouse.

**Extended Figure 10.**
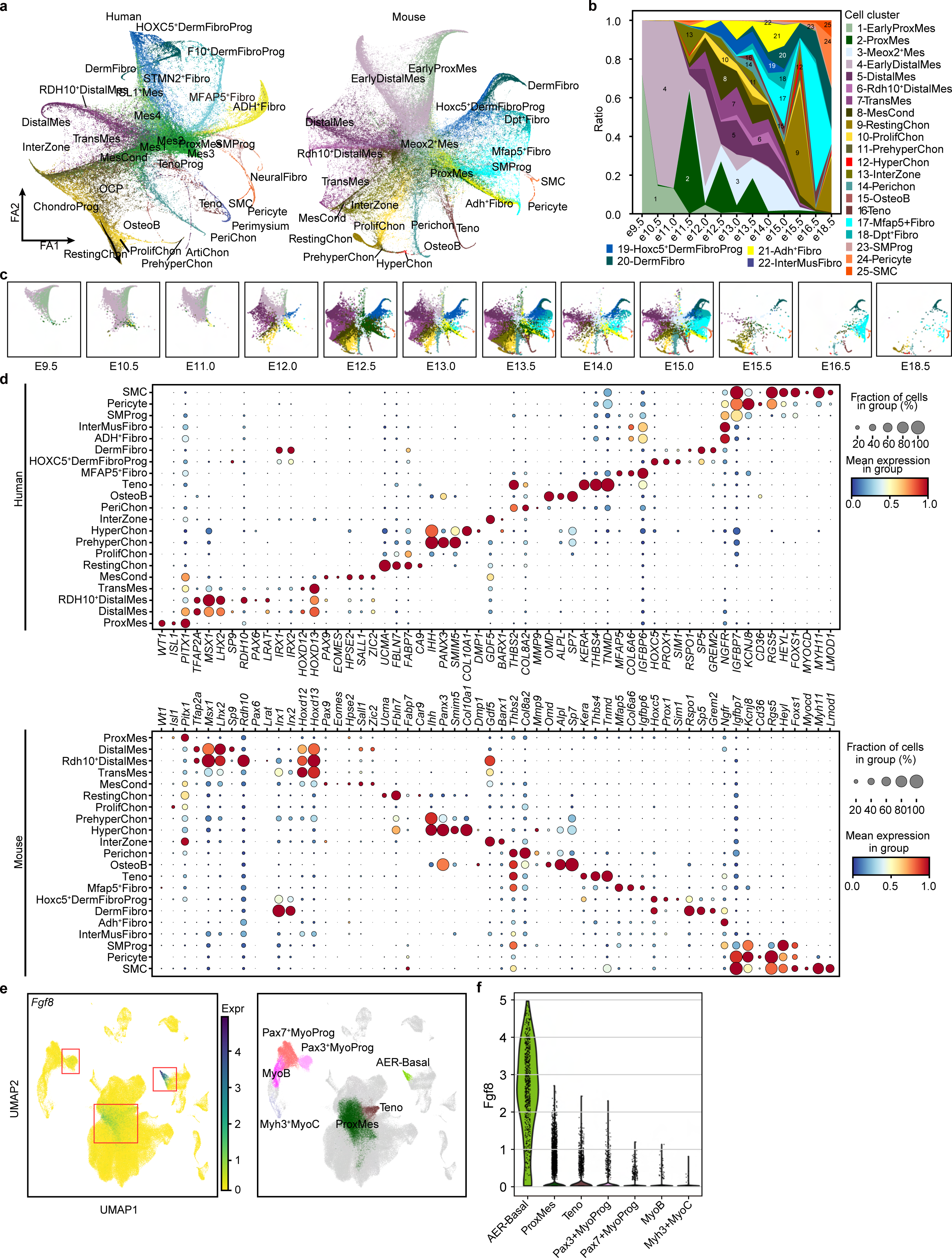
The LPM lineage in human and mouse. **a**, Force-directed graph of human and mouse LPM-derived cells, colored by cell clusters. **b**, Stacked bar chart showing the fraction of mouse cell clusters per time point, followed by the color code of mouse cell clusters in a. **c**, Force-directed graph of mouse cells per time point, coloured by cell cluster in a. **d**, Dotplot of selected transcription factors expressed in human and mouse.

## Data Availability

All of our newly generated raw data are publicly available on ArrayExpress (mouse scRNA-seq, E-MTAB-10514; human Visium, E-MTAB-10367; human scRNA-seq, E-MTAB-8813). Previously published raw data can be found from ENCODE portal (ENCSR713GIS) and GEO (GSE137335 and GSE142425). Processed data and be downloaded and visualized at our data portal (https://limb-dev.cellgeni.sanger.ac.uk/).

## Code availability

All in-house codes can be found on github (https://github.com/Teichlab/limbcellatlas/).

## Author contributions

S.A.T. and H.Z. supervised the project; S.A.T. initiated and designed the project; X.H. and Y.F. carried out human tissue collection; B.W. carried out mouse tissue collection; B.W.,L.M., L.B., R.E., and E.F. performed scRNA-seq; E.T. performed Visium spatial experiments; S.W.,K.R., and E.T did in situ staining and functional experiments;AC and RB performed light sheet fluorescence microscopy. H.Y., C.L. and H.Z. provided experimental support. B.Z.,P.H and J.E.L. analysed sequencing data and generated figures. V.K., K.P., M.P. and N.Y. provided computational support. M.S. and B.J.W.contributed to interpretation of the results. B.Z.,P.H, J.E.L., S.W., H.Z. and S.A.T. wrote the manuscript. All authors contributed to the discussion and editing of the manuscript.

## Supporting information

Table 1 - Marker Genes

Table 2 - Spatial genes in foot plate

Table 3 - Ortholog phenotypes in mouse and human

Table 4 - SCENIC scores

Table 5 - Sample metadata

## Acknowledgements

We thank Jana Lalakova for illustrating human limb syndromes. We thank Ken To for proofreading the manuscript. We thank Matt Thomson’s lab for help with mouse 10X loading. We thank members of the Teichmann lab, Zhang lab, Marioni lab, Haniffa lab and Behjati lab for discussion and feedback. We thank Olivier Pourquié for feedback on muscle development.

This work was supported by the Wellcome Trust Grant 206194, 108413/A/15/D and 211276/Z/18/Z, as well as the National Key Research and Development Program (grant 2019YFA0801703), National Natural Science Foundation of China (grant 31871370), Science and Technology Program of Guangzhou (grant 202002030429), and Advanced Medical Technology Center, The First Affiliated Hospital, Zhongshan School of Medicine, Sun Yat-sen University (to H.Z.); China Postdoctoral Science Foundation (grant 2021M700936), and Natural Science Foundation of Guangdong (grant 2019A1515011342) (to S.W.). P.H holds a non-stipendiary research fellowship at St Edmund’s College, University of Cambridge. J.E.L is funded by the Wellcome Trust under the clinical PhD programme (grant 222902/Z/21/Z) and supported by Darwin College, Cambridge through a Geoffrey Fisk Studentship. A.C is funded by the Inserm cross-cutting program HuDeCA 2018. R.B was recipient of a fellowship from the “Fondation pour la Recherche Médicale” (FRM).

## References

1. Chevallier, A., Kieny, M. & Mauger, A. Limb-somite relationship: origin of the limb musculature. J. Embryol. Exp. Morphol. 41, 245–258 (1977).

2. Christ, B., Jacob, H. J. & Jacob, M. Experimental analysis of the origin of the wing musculature in avian embryos. Anat. Embryol. 150, 171–186 (1977).

3. Tabin, C. & Wolpert, L. Rethinking the proximodistal axis of the vertebrate limb in the molecular era. Genes Dev. 21, 1433–1442 (2007).

4. Zuniga, A. Next generation limb development and evolution: old questions, new perspectives. Development 142, 3810–3820 (2015).

5. Hawkins, M. B., Henke, K. & Harris, M. P. Latent developmental potential to form limb-like skeletal structures in zebrafish. Cell 184, 899–911.e13 (2021).

6. Lopez-Rios, J. The many lives of SHH in limb development and evolution. Semin. Cell Dev. Biol. 49, 116–124 (2016).

7. McQueen, C. & Towers, M. Establishing the pattern of the vertebrate limb. Development 147, dev177956 (2020).

8. Petit, F., Sears, K. E. & Ahituv, N. Limb development: a paradigm of gene regulation. Nat. Rev. Genet. 18, 245–258 (2017).

9. Moore, K. L., Persaud, T. V. N. & Torchia, M. G. The Developing Human - E-Book: Clinically Oriented Embryology. (Elsevier Health Sciences, 2018).

10. Wilkie, A. O. M. Why study human limb malformations? J. Anat. 202, 27–35 (2003).

11. Kildisiute, G. et al. Tumor to normal single-cell mRNA comparisons reveal a pan-neuroblastoma cancer cell. Sci Adv 7, (2021).

12. Elmentaite, R. et al. Cells of the human intestinal tract mapped across space and time. Nature 597, 250–255 (2021).

13. Elmentaite, R. et al. Single-Cell Sequencing of Developing Human Gut Reveals Transcriptional Links to Childhood Crohn’s Disease. Dev. Cell 55, 771–783.e5 (2020).

14. Garcia-Alonso, L. et al. Mapping the temporal and spatial dynamics of the human endometrium in vivo and in vitro. bioRxiv 2021.01.02.425073 (2021) doi:10.1101/2021.01.02.425073.

15. Jardine, L. et al. Blood and immune development in human fetal bone marrow and Down syndrome. Nature 598, 327–331 (2021).

16. He, P. et al. A human fetal lung cell atlas uncovers proximal-distal gradients of differentiation and key regulators of epithelial fates. Cell 185, 4841–4860.e25 (2022).

17. Behjati, S., Lindsay, S., Teichmann, S. A. & Haniffa, M. Mapping human development at single-cell resolution. Development 145, (2018).

18. Haniffa, M. et al. A roadmap for the Human Developmental Cell Atlas. Nature 597, 196–205 (2021).

19. Xi, H. et al. A Human Skeletal Muscle Atlas Identifies the Trajectories of Stem and Progenitor Cells across Development and from Human Pluripotent Stem Cells. Cell Stem Cell 27, 158–176.e10 (2020).

20. Kleshchevnikov, V. et al. Cell2location maps fine-grained cell types in spatial transcriptomics. Nat. Biotechnol. 1–11 (2022).

21. Satoda, M., Pierpont, M. E., Diaz, G. A., Bornemeier, R. A. & Gelb, B. D. Char syndrome, an inherited disorder with patent ductus arteriosus, maps to chromosome 6p12-p21. Circulation 99, (1999).

22. Cunningham, T. J., Chatzi, C., Sandell, L. L., Trainor, P. A. & Duester, G. Rdh10 Mutants Deficient in Limb Field Retinoic Acid Signaling Exhibit Normal Limb Patterning but Display Interdigital Webbing. Dev. Dyn. 240, 1142 (2011).

23. Díaz-Hernández, M. E., Bustamante, M., Galván-Hernández, C. I. & Chimal-Monroy, J. Irx1 and Irx2 Are Coordinately Expressed and Regulated by Retinoic Acid, TGFβ and FGF Signaling during Chick Hindlimb Development. PLoS One 8, e58549 (2013).

24. Zülch, A., Becker, M. B. & Gruss, P. Expression pattern of Irx1 and Irx2 during mouse digit development. Mech. Dev. 106, (2001).

25. Richardson, R. J. et al. Periderm prevents pathological epithelial adhesions during embryogenesis. J. Clin. Invest. 124, 3891–3900 (2014).

26. Saunders, J. W., Jr. The proximo-distal sequence of origin of the parts of the chick wing and the role of the ectoderm. J. Exp. Zool. 108, 363–403 (1948).

27. Towers, M. & Tickle, C. Growing models of vertebrate limb development. Development 136, (2009).

28. Riddle, R. D., Johnson, R. L., Laufer, E. & Tabin, C. Sonic hedgehog mediates the polarizing activity of the ZPA. Cell 75, 1401–1416 (1993).

29. Zúñiga, A., Haramis, A. P., McMahon, A. P. & Zeller, R. Signal relay by BMP antagonism controls the SHH/FGF4 feedback loop in vertebrate limb buds. Nature 401, 598–602 (1999).

30. Towers, M., Mahood, R., Yin, Y. & Tickle, C. Integration of growth and specification in chick wing digit-patterning. Nature 452, 882–886 (2008).

31. Ballard, K. J. & Holt, S. J. Cytological and cytochemical studies on cell death and digestion in the foetal rat foot: the role of macrophages and hydrolytic enzymes. J. Cell Sci. 3, 245–262 (1968).

32. Hernández-Martínez, R., Castro-Obregón, S. & Covarrubias, L. Progressive interdigital cell death: regulation by the antagonistic interaction between fibroblast growth factor 8 and retinoic acid. Development 136, (2009).

33. Salas-Vidal, E., Valencia, C. & Covarrubias, L. Differential tissue growth and patterns of cell death in mouse limb autopod morphogenesis. Dev. Dyn. 220, (2001).

34. Capdevila, J., Tsukui, T., Rodríquez, E. C., Zappavigna, V. & Jc, I. B. Control of vertebrate limb outgrowth by the proximal factor Meis2 and distal antagonism of BMPs by Gremlin. Mol. Cell 4, (1999).

35. Penkov, D. et al. Analysis of the DNA-binding profile and function of TALE homeoproteins reveals their specialization and specific interactions with Hox genes/proteins. Cell Rep. 3, (2013).

36. Delgado, I. et al. Control of mouse limb initiation and antero-posterior patterning by Meis transcription factors. Nat. Commun. 12, 1–13 (2021).

37. Formation of Proximal and Anterior Limb Skeleton Requires Early Function of Irx3 and Irx5 and Is Negatively Regulated by Shh Signaling. Dev. Cell 29, 233– 240 (2014).

38. Yamaguchi, T. P., Bradley, A., McMahon, A. P. & Jones, S. A Wnt5a pathway underlies outgrowth of multiple structures in the vertebrate embryo. Development 126, (1999).

39. Guo-hao Lin, L.Z.. Apical ectodermal ridge regulates three principal axes of the developing limb. J. Zhejiang Univ. Sci. B 21, 757 (2020).

40. Mao, J., McGlinn, E., Huang, P., Tabin, C. J. & McMahon, A. P. Fgf-dependent Etv4/5 activity is required for posterior restriction of Sonic Hedgehog and promoting outgrowth of the vertebrate limb. Dev. Cell 16, (2009).

41. Kawakami, Y. et al. Sall genes regulate region-specific morphogenesis in the mouse limb by modulating Hox activities. Development 136, 585 (2009).

42. Fernandez-Teran, M. et al. Role of dHAND in the anterior-posterior polarization of the limb bud: implications for the Sonic hedgehog pathway. Development 127, 2133–2142 (2000).

43. McGlinn, E. et al. Pax9 and Jagged1 act downstream of Gli3 in vertebrate limb development. Mech. Dev. 122, (2005).

44. Kuijper, S. et al. Function and regulation of Alx4 in limb development: complex genetic interactions with Gli3 and Shh. Dev. Biol. 285, (2005).

45. Quinn, M. E., Haaning, A. & Ware, S. M. Preaxial polydactyly caused by Gli3 haploinsufficiency is rescued by Zic3 loss of function in mice. Hum. Mol. Genet. 21, (2012).

46. Galli, A. et al. Distinct roles of Hand2 in initiating polarity and posterior Shh expression during the onset of mouse limb bud development. PLoS Genet. 6, e1000901 (2010).

47. Fuse, N. et al. Sonic hedgehog protein signals not as a hydrolytic enzyme but as an apparent ligand for Patched. Proc. Natl. Acad. Sci. U. S. A. 96, 10992–10999 (1999).

48. Huangfu, D. & Anderson, K. V. Signaling from Smo to Ci/Gli: conservation and divergence of Hedgehog pathways from Drosophila to vertebrates. Development 133, 3–14 (2006).

49. The role of Hox genes during vertebrate limb development. Curr. Opin. Genet. Dev. 17, 359–366 (2007).

50. Tarchini, B. & Duboule, D. Control of Hoxd genes’ collinearity during early limb development. Dev. Cell 10, 93–103 (2006).

51. Dollé, P., Izpisúa-Belmonte, J. C., Falkenstein, H., Renucci, A. & Duboule, D. Coordinate expression of the murine Hox-5 complex homoeobox-containing genes during limb pattern formation. Nature 342, 767–772 (1989).

52. Kherdjemil, Y. et al. Evolution of Hoxa11 regulation in vertebrates is linked to the pentadactyl state. Nature 539, (2016).

53. McCulloch, D. R. et al. ADAMTS metalloproteases generate active versican fragments that regulate interdigital web regression. Dev. Cell 17, 687 (2009).

54. Kaltcheva, M. M., Anderson, M. J., Harfe, B. D. & Lewandoski, M. BMPs are direct triggers of interdigital programmed cell death. Dev. Biol. 411, 266 (2016).

55. Díaz-Hernández, M. E., Rios-Flores, A. J., Abarca-Buis, R. F., Bustamante, M. & Chimal-Monroy, J. Molecular Control of Interdigital Cell Death and Cell Differentiation by Retinoic Acid during Digit Development. Journal of Developmental Biology 2, 138–157 (2014).

56. Spagnoli, A. et al. TGF-β signaling is essential for joint morphogenesis. J. Cell Biol. 177, 1105 (2007).

57. Mokuda, S. et al. Wwp2 maintains cartilage homeostasis through regulation of Adamts5. Nat. Commun. 10, 1–13 (2019).

58. Wu, C.-L. et al. Single cell transcriptomic analysis of human pluripotent stem cell chondrogenesis. Nat. Commun. 12, 1–18 (2021).

59. Zhou, T. et al. Piezo1/2 mediate mechanotransduction essential for bone formation through concerted activation of NFAT-YAP1-ß-catenin. Elife 9, (2020).

60. Lorda-Diez, C. I., Torre-Pérez, N., García-Porrero, J. A., Hurle, J. M. & Montero, J. A. Expression of Id2 in the developing limb is associated with zones of active BMP signaling and marks the regions of growth and differentiation of the developing digits. Int. J. Dev. Biol. 53, (2009).

61. McGlinn, E. et al. Expression of the NET family member Zfp503 is regulated by hedgehog and BMP signaling in the limb. Dev. Dyn. 237, 1172–1182 (2008).

62. Ma, S., Charron, J. & Erikson, R. L. Role of Plk2 (Snk) in Mouse Development and Cell Proliferation. Mol. Cell. Biol. 23, 6936 (2003).

63. Sasahira, T., Kurihara, M., Nakashima, C., Kirita, T. & Kuniyasu, H. LEM domain containing 1 promotes oral squamous cell carcinoma invasion and endothelial transmigration. Br. J. Cancer 115, 52–58 (2016).

64. Falardeau, F., Camurri, M. V. & Campeau, P. M. Genomic approaches to diagnose rare bone disorders. Bone 102, 5–14 (2017).

65. Cooks, R. G., Hertz, M., Katznelson, M. B. & Goodman, R. M. A new nail dysplasia syndrome with onychonychia and absence and/or hypoplasia of distal phalanges. Clin. Genet. 27, (1985).

66. Temtamy, S. A. & Aglan, M. S. Brachydactyly. Orphanet J. Rare Dis. 3, 1–16 (2008).

67. Bahubali D. Gane, P.N. Split-hand/feet malformation: A rare syndrome. Journal of Family Medicine and Primary Care 5, 168 (2016).

68. Marcelis, C. L. M. & de Brouwer, A. P. M. Feingold Syndrome 1. In GeneReviews® [Internet] (University of Washington, Seattle, 2019).

69. Mutations in BMP4 Cause Eye, Brain, and Digit Developmental Anomalies: Overlap between the BMP4 and Hedgehog Signaling Pathways. Am. J. Hum. Genet. 82, 304–319 (2008).

70. El Ghouzzi, V., et al. Saethre-Chotzen mutations cause TWIST protein degradation or impaired nuclear location. Hum. Mol. Genet. 9, (2000).

71. Eiken, M., Prag, J., Petersen, K. E. & Kaufmann, H. J. A new familial skeletal dysplasia with severely retarded ossification and abnormal modeling of bones especially of the epiphyses, the hands, and feet. Eur. J. Pediatr. 141, (1984).

72. Fan, Y. et al. Parathyroid hormone 1 receptor is essential to induce FGF23 production and maintain systemic mineral ion homeostasis. The FASEB Journal 30, 428 (2016).

73. 73Yi, S. E., Daluiski, A., Pederson, R., Rosen, V. & Lyons, K. M. The type I BMP receptor BMPRIB is required for chondrogenesis in the mouse limb. Development vol. 127 621–630 Preprint at https://doi.org/10.1242/dev.127.3.621 (2000).

74. Stafford, D. A., Brunet, L. J., Khokha, M. K., Economides, A. N. & Harland, R. M. Cooperative activity of noggin and gremlin 1 in axial skeleton development. Development 138, 1005–1014 (2011).

75. Brunet, L. J., McMahon, J. A., McMahon, A. P. & Harland, R. M. Noggin, Cartilage Morphogenesis, and Joint Formation in the Mammalian Skeleton. Science vol. 280 1455–1457 Preprint at https://doi.org/10.1126/science.280.5368.1455 (1998).

76. Melkoniemi, M. et al. Autosomal recessive disorder otospondylomegaepiphyseal dysplasia is associated with loss-of-function mutations in the COL11A2 gene. Am. J. Hum. Genet. 66, 368–377 (2000).

77. Li, S. W. et al. Targeted disruption of Col11a2 produces a mild cartilage phenotype in transgenic mice: comparison with the human disorder otospondylomegaepiphyseal dysplasia (OSMED). Dev. Dyn. 222, 141–152 (2001).

78. Bi, W. et al. Haploinsufficiency of Sox9 results in defective cartilage primordia and premature skeletal mineralization. Proc. Natl. Acad. Sci. U. S. A. 98, 6698– 6703 (2001).

79. Shazeeb, M. S. et al. Skeletal Characterization of the Fgfr3 Mouse Model of Achondroplasia Using Micro-CT and MRI Volumetric Imaging. Scientific Reports vol. 8 Preprint at https://doi.org/10.1038/s41598-017-18801-0 (2018).

80. Carey, J. C., Cassidy, S. B., Battaglia, A. & Viskochil, D. Cassidy and Allanson’s Management of Genetic Syndromes. (John Wiley & Sons, 2021).

81. Sowińska-Seidler, A., Socha, M. & Jamsheer, A. Split-hand/foot malformation-molecular cause and implications in genetic counseling. J. Appl. Genet. 55, 105–115 (2014).

82. Merlo, G. R. et al. Mouse model of split hand/foot malformation type I. Genesis 33, 97–101 (2002).

83. Celli, J., van Bokhoven, H. & Brunner, H. G. Feingold syndrome: Clinical review and genetic mapping. American Journal of Medical Genetics vol. 122A 294–300 Preprint at https://doi.org/10.1002/ajmg.a.20471 (2003).

84. Sawai, S. et al. Defects of embryonic organogenesis resulting from targeted disruption of the N-myc gene in the mouse. Development 117, 1445–1455 (1993).

85. Selever, J., Liu, W., Lu, M.-F., Behringer, R. R. & Martin, J. F. Bmp4 in limb bud mesoderm regulates digit pattern by controlling AER development. Developmental Biology vol. 276 268–279 Preprint at https://doi.org/10.1016/j.ydbio.2004.08.024 (2004).

86. Gripp, K. W., Zackai, E. H. & Stolle, C. A. Mutations in the human TWIST gene. Hum. Mutat. 15, 479 (2000).

87. Bialek, P. et al. A Twist Code Determines the Onset of Osteoblast Differentiation. Developmental Cell vol. 6 423–435 Preprint at https://doi.org/10.1016/s1534-5807(04)00058-9 (2004).

88. Moore, A. W. et al. YAC transgenic analysis reveals Wilms’ tumour 1 gene activity in the proliferating coelomic epithelium, developing diaphragm and limb. Mech. Dev. 79, (1998).

89. Distinct populations within Isl1 lineages contribute to appendicular and facial skeletogenesis through the β-catenin pathway. Dev. Biol. 387, 37–48 (2014).

90. Dunwoodie, S. L., Rodriguez, T. A. & Beddington, R. S. Msg1 and Mrg1, founding members of a gene family, show distinct patterns of gene expression during mouse embryogenesis. Mech. Dev. 72, 27–40 (1998).

91. Antin, P. B., Pier, M., Sesepasara, T., Yatskievych, T. A. & Darnell, D. K. Embryonic Expression of the Chicken Krüppel-like (KLF) Transcription Factor Gene Family. Dev. Dyn. 239, 1879 (2010).

92. Hu, W. Y. et al. Isolation and functional interrogation of adult human prostate epithelial stem cells at single cell resolution. Stem Cell Res. 23, (2017).

93. Aibar, S. et al. SCENIC: single-cell regulatory network inference and clustering. Nat. Methods 14, 1083–1086 (2017).

94. Hudson, D. T. et al. Gene expression analysis of the Xenopus laevis early limb bud proximodistal axis. Dev. Dyn. 251, (2022).

95. Sheng, G. & Stern, C. D. Gata2 and Gata3: novel markers for early embryonic polarity and for non-neural ectoderm in the chick embryo. Mech. Dev. 87, (1999).

96. Tzchori, I. et al. LIM homeobox transcription factors integrate signaling events that control three-dimensional limb patterning and growth. Development 136, 1375–1385 (2009).

97. Bensoussan-Trigano, V., Lallemand, Y., Saint, C. C. & Robert, B. Msx1 and Msx2 in limb mesenchyme modulate digit number and identity. Dev. Dyn. 240, (2011).

98. Fromental-Ramain, C. et al. Hoxa-13 and Hoxd-13 play a crucial role in the patterning of the limb autopod. Development 122, (1996).

99. Arostegui, M., Scott, R. W., Böse, K. & Underhill, T. M. Cellular taxonomy of Hic1+ mesenchymal progenitor derivatives in the limb: from embryo to adult. Nat. Commun. 13, 1–20 (2022).

100. Liu, C. F. & Lefebvre, V. The transcription factors SOX9 and SOX5/SOX6 cooperate genome-wide through super-enhancers to drive chondrogenesis. Nucleic Acids Res. 43, (2015).

101. Kawato, Y. et al. Nkx3.2 promotes primary chondrogenic differentiation by upregulating Col2a1 transcription. PLoS One 7, (2012).

102. Hong, E., Di Cesare, P. E. & Haudenschild, D. R. Role of c-Maf in Chondrocyte Differentiation: A Review. Cartilage 2, 27 (2011).

103. Venugopalan, S. R. et al. Hierarchical interactions of homeodomain and forkhead transcription factors in regulating odontogenic gene expression. J. Biol. Chem. 286, 21372–21383 (2011).

104. Dreher, S. I., Fischer, J., Walker, T., Diederichs, S. & Richter, W. Significance of MEF2C and RUNX3 Regulation for Endochondral Differentiation of Human Mesenchymal Progenitor Cells. Front Cell Dev Biol 8, 81 (2020).

105. Ghoul-Mazgar, S. et al. Expression pattern of Dlx3 during cell differentiation in mineralized tissues. Bone 37, 799–809 (2005).

106. Yan, J. et al. Smad4 deficiency impairs chondrocyte hypertrophy via the Runx2 transcription factor in mouse skeletal development. J. Biol. Chem. 293, 9162– 9175 (2018).

107. Zhang, J. et al. Roles of SATB2 in Osteogenic Differentiation and Bone Regeneration. Tissue Eng. Part A 17, 1767 (2011).

108. Iwamoto, M. et al. Transcription factor ERG and joint and articular cartilage formation during mouse limb and spine skeletogenesis. Dev. Biol. 305, (2007).

109. Zhao, H. et al. Foxp1/2/4 regulate endochondral ossification as a suppresser complex. Dev. Biol. 398, 242 (2015).

110. Pitsillides, A. A. & Beier, F. Keep your Sox on, chondrocytes! Nat. Rev. Rheumatol. 17, 383–384 (2021).

111. Henry, S. P., Liang, S., Akdemir, K. C. & de Crombrugghe, B. The postnatal role of Sox9 in cartilage. J. Bone Miner. Res. 27, 2511–2525 (2012).

112. Blitz, E., Sharir, A., Akiyama, H. & Zelzer, E. Tendon-bone attachment unit is formed modularly by a distinct pool of Scx-and Sox9-positive progenitors. Development 140, 2680–2690 (2013).

113. Yang, F. & Richardson, D. W. Comparative Analysis of Tenogenic Gene Expression in Tenocyte-Derived Induced Pluripotent Stem Cells and Bone Marrow-Derived Mesenchymal Stem Cells in Response to Biochemical and Biomechanical Stimuli. Stem Cells Int. 2021, (2021).

114. Rossi, G. et al. Nfix Regulates Temporal Progression of Muscle Regeneration through Modulation of Myostatin Expression. Cell Rep. 14, (2016).

115. Hoi, C. S. L. et al. Runx1 Directly Promotes Proliferation of Hair Follicle Stem Cells and Epithelial Tumor Formation in Mouse Skin. Mol. Cell. Biol. 30, 2518 (2010).

116. Li, L. et al. TFAP2C-and p63-Dependent Networks Sequentially Rearrange Chromatin Landscapes to Drive Human Epidermal Lineage Commitment. Cell Stem Cell 24, (2019).

117. Lepore, J. J., Cappola, T. P., Mericko, P. A., Morrisey, E. E. & Parmacek, M. S. GATA-6 regulates genes promoting synthetic functions in vascular smooth muscle cells. Arterioscler. Thromb. Vasc. Biol. 25, (2005).

118. Xie, Z. et al. Smooth-muscle BMAL1 participates in blood pressure circadian rhythm regulation. J. Clin. Invest. 125, (2015).

119. Buckingham, M. & Rigby, P. W. J. Gene regulatory networks and transcriptional mechanisms that control myogenesis. Dev. Cell 28, 225–238 (2014).

120. Ontell, M. & Kozeka, K. The organogenesis of murine striated muscle: a cytoarchitectural study. Am. J. Anat. 171, 133–148 (1984).

121. Buckingham, M. et al. The formation of skeletal muscle: from somite to limb. J. Anat. 202, 59–68 (2003).

122. Hutcheson, D. A., Zhao, J., Merrell, A. & Haldar, M. Embryonic and fetal limb myogenic cells are derived from developmentally distinct progenitors and have different requirements for β-catenin. Genes (2009).

123. Singh, A. J. et al. FACS-Seq analysis of Pax3-derived cells identifies non-myogenic lineages in the embryonic forelimb. Sci. Rep. 8, 7670 (2018).

124. Benezra, R., Davis, R. L., Lockshon, D., Turner, D. L. & Weintraub, H. The protein Id: a negative regulator of helix-loop-helix DNA binding proteins. Cell 61, 49–59 (1990).

125. Roschger, C. & Cabrele, C. The Id-protein family in developmental and cancer-associated pathways. Cell Commun. Signal. 15, 7 (2017).

126. Muriel, J. M. et al. Keratin 18 is an integral part of the intermediate filament network in murine skeletal muscle. Am. J. Physiol. Cell Physiol. 318, (2020).

127. Hernandez-Torres, F., Rodríguez-Outeiriño, L., Franco, D. & Aranega, A. E. Pitx2 in Embryonic and Adult Myogenesis. Front Cell Dev Biol 5, 46 (2017). 128.

128. Lee, H., Habas, R. & Abate-Shen, C. MSX1 cooperates with histone H1b for inhibition of transcription and myogenesis. Science 304, (2004).

129. Relaix, F. et al. Six homeoproteins directly activate Myod expression in the gene regulatory networks that control early myogenesis. PLoS Genet. 9, (2013).

130. Meech, R. et al. Barx2 is Expressed in Satellite Cells and is Required for Normal Muscle Growth and Regeneration. Stem Cells 30, 253 (2012).

131. Zappia, M. P. & Frolov, M. V. E2F function in muscle growth is necessary and sufficient for viability in Drosophila. Nat. Commun. 7, (2016).

132. Lazure, F. et al. Myf6/MRF4 is a myogenic niche regulator required for the maintenance of the muscle stem cell pool. EMBO Rep. 21, (2020).

133. Lu, J., Webb, R., Richardson, J. A. & Olson, E. N. MyoR: A muscle-restricted basic helix–loop–helix transcription factor that antagonizes the actions of MyoD. Proc. Natl. Acad. Sci. U. S. A. 96, 552–557 (1999).

134. MacQuarrie, K. L., Yao, Z., Fong, A. P. & Tapscott, S. J. Genome-wide binding of the basic helix-loop-helix myogenic inhibitor musculin has substantial overlap with MyoD: implications for buffering activity. Skelet. Muscle 3, 26 (2013).

135. Buas, M. F., Kabak, S. & Kadesch, T. Inhibition of myogenesis by Notch: evidence for multiple pathways. J. Cell. Physiol. 218, 84–93 (2009).

136. Chiu, Y.-K. et al. Transcription factor ABF-1 suppresses plasma cell differentiation but facilitates memory B cell formation. J. Immunol. 193, 2207– 2217 (2014).

137. Massari, M. E. et al. Characterization of ABF-1, a novel basic helix-loop-helix transcription factor expressed in activated B lymphocytes. Mol. Cell. Biol. 18, 3130–3139 (1998).

138. Efremova, M., Vento-Tormo, M., Teichmann, S. A. & Vento-Tormo, R. CellPhoneDB: inferring cell-cell communication from combined expression of multi-subunit ligand-receptor complexes. Nat. Protoc. 15, 1484–1506 (2020).

139. Vento-Tormo, R. et al. Single-cell reconstruction of the early maternal–fetal interface in humans. Nature 563, 347–353 (2018).

140. Kawakami, Y. et al. Identification of chick frizzled-10 expressed in the developing limb and the central nervous system. Mech. Dev. 91, (2000).

141. Nunnally, A. P. & Parr, B. A. Analysis of Fz10 expression in mouse embryos. Dev. Genes Evol. 214, (2004).

142. Sarem, M., Otto, O., Tanaka, S. & Shastri, V. P. Cell number in mesenchymal stem cell aggregates dictates cell stiffness and chondrogenesis. Stem Cell Res. Ther. 10, 1–18 (2019).

143. Schroeter, E. H., Kisslinger, J. A. & Kopan, R. Notch-1 signalling requires ligand-induced proteolytic release of intracellular domain. Nature 393, 382–386 (1998).

144. D’Souza, B., Miyamoto, A. & Weinmaster, G. The many facets of Notch ligands. Oncogene 27, 5148–5167 (2008).

145. Canonical Notch ligands and Fringes have distinct effects on NOTCH1 and NOTCH2. J. Biol. Chem. 295, 14710–14722 (2020).

146. Crosnier, C. et al. JAGGED1 gene expression during human embryogenesis elucidates the wide phenotypic spectrum of Alagille syndrome. Hepatology 32, (2000).

147. Mašek, J. & Andersson, E. R. The developmental biology of genetic Notch disorders. Development 144, 1743–1763 (2017).

148. Turnpenny, P. D. & Ellard, S. Alagille syndrome: pathogenesis, diagnosis and management. Eur. J. Hum. Genet. 20, 251–257 (2011).

149. Xu, X. et al. Fibroblast growth factor receptor 2 (FGFR2)-mediated reciprocal regulation loop between FGF8 and FGF10 is essential for limb induction. Development (1998).

150. Agha, E. E., et al. Characterization of a novel Fibroblast growth factor 10 (Fgf10) knock-in mouse line to target mesenchymal progenitors during embryonic development. Pneumologie vol. 66 Preprint at https://doi.org/10.1055/s-0032-1315504 (2012).

151. Azoury, S. C., Reddy, S., Shukla, V. & Deng, C.-X. Fibroblast Growth Factor Receptor 2 (FGFR2) Mutation Related Syndromic Craniosynostosis. Int. J. Biol. Sci. 13, 1479 (2017).

152. He, P. et al. The changing mouse embryo transcriptome at whole tissue and single-cell resolution. Nature 583, 760–767 (2020).

153. Kelly, N. H., Huynh, N. P. T. & Guilak, F. Single cell RNA-sequencing reveals cellular heterogeneity and trajectories of lineage specification during murine embryonic limb development. Matrix Biology vol. 89 1–10 Preprint at https://doi.org/10.1016/j.matbio.2019.12.004 (2020).

154. Allou, L. et al. Non-coding deletions identify Maenli lncRNA as a limb-specific En1 regulator. Nature 592, 93–98 (2021).

155. Jain, M. S. et al. MultiMAP: dimensionality reduction and integration of multimodal data. Genome Biol. 22, 1–26 (2021).

156. Gilson, H. et al. Follistatin induces muscle hypertrophy through satellite cell proliferation and inhibition of both myostatin and activin. Am. J. Physiol. Endocrinol. Metab. 297, (2009).

157. Gao, H. et al. UCHL1 regulates muscle fibers and mTORC1 activity in skeletal muscle. Life Sci. 233, 116699 (2019).

158. Purushothaman, S., Elewa, A. & Seifert, A. W. Fgf-signaling is compartmentalized within the mesenchyme and controls proliferation during salamander limb development. (2019) doi:10.7554/eLife.48507.

159. Popescu, D. M. et al. Decoding human fetal liver haematopoiesis. Nature 574, (2019).

160. Wolock, S. L., Lopez, R. & Klein, A. M. Scrublet: Computational Identification of Cell Doublets in Single-Cell Transcriptomic Data. Cell Syst 8, 281–291.e9 (2019).

161. Pedregosa, F. et al. Scikit-learn: Machine Learning in Python. J. Mach. Learn. Res. 12, 2825–2830 (2011).

162. Schindelin, J., et al. Fiji: an open-source platform for biological-image analysis. Nature Methods vol. 9 676–682 Preprint at https://doi.org/10.1038/nmeth.2019 (2012).

163. Bergen, V., Lange, M., Peidli, S., Wolf, F. A. & Theis, F. J. Generalizing RNA velocity to transient cell states through dynamical modeling. Nat. Biotechnol. 38, 1408–1414 (2020).

164. Van de Sande, B. et al. A scalable SCENIC workflow for single-cell gene regulatory network analysis. Nat. Protoc. 15, 2247–2276 (2020).

165. Wang, S., et al. Muscle Stem Cell Immunostaining. Curr. Protoc. Mouse Biol. 8, e47 (2018).

166. Belle, M. et al. Tridimensional Visualization and Analysis of Early Human Development. Cell 169, 161–173.e12 (2017).

167. Lapan, A. D. & Gussoni, E. Isolation and characterization of human fetal myoblasts. Methods Mol. Biol. 798, 3–19 (2012).

